# Spatial atlas highlights contribution of *C. difficile* in early-stage colorectal cancer

**DOI:** 10.64898/2026.06.18.733268

**Authors:** Nicholas O. Markham, Julia L. Drewes, Emily H. Green, William E. Ball, Hannah M. Lunnemann, Alan J. Simmons, Harsimran Kaur, Jessica Queen, Abby L. Geis, Maryam Pourmaleki, Bryan R. Helm, Erik Storrs, Marisol Ramirez-Solano, Xiang Li, Zhicheng Ma, Frank Revetta, Annika Windon, M. Kay Washington, Jane W. Wanyiri, Thevambiga Iyadorai, April C. Roslani, Jamuna Vadivelu, Jonathan Coggin, Leslie M. Meenderink, D. Borden Lacy, Christina Curtis, Siyuan Ma, Li Ding, Qi Liu, Martha J. Shrubsole, Ken S. Lau, Cynthia L. Sears, Robert J. Coffey

**Affiliations:** Department of Veterans Affairs, Tennessee Valley Healthcare System, Nashville, TN; Epithelial Biology Center, VUMC, Nashville, TN; Vanderbilt-Ingram Cancer Center, VUMC, Nashville, TN; Department of Medicine, Division of Gastroenterology, Hepatology, and Nutrition, VUMC, Nashville, TN; Department of Pathology, Microbiology, and Immunology, VUMC, Nashville, TN; Vanderbilt Institute for Infection, Inflammation, and Immunology, VUMC, Nashville, TN; Division of Infectious Diseases, Department of Medicine, Johns Hopkins University School of Medicine, Baltimore, MD; Department of Oncology, Johns Hopkins University School of Medicine, Baltimore, MD; Department of Cell and Developmental Biology, Vanderbilt University, Nashville, TN; Chemical and Physical Biology Program, Vanderbilt University, Nashville, TN; Stanford Cancer Institute, Stanford University School of Medicine, Stanford, CA; Department of Biostatistics, VUMC, Nashville, TN; Department of Medicine, Washington University in St. Louis, St. Louis, MO; McDonnell Genome Institute, Washington University in St. Louis, St. Louis, MO; Department of Medicine, Division of Oncology, Stanford University School of Medicine, Stanford, CA; Weill Cornell Medical College and New York Presbyterian Hospital, New York City, NY; Department of Molecular Microbiology and Immunology, Bloomberg School of Public Health, Johns Hopkins University, Baltimore, MD; Faculty of Medicine, Universiti Malaya, Kuala Lumpur, Malaysia; Chan Zuckerberg Biohub, San Francisco, CA; Siteman Cancer Center, Washington University in St. Louis, St. Louis, MO; Department of Genetics, Washington University in St Louis, St Louis, MO; Department of Medicine, Division of Epidemiology, VUMC, Nashville, TN; Center for Computational Systems Biology, Vanderbilt University, Nashville TN; Department of Surgery, Vanderbilt University Medical Center, Nashville TN

**Keywords:** Colorectal cancer, microbiome, spatial transcriptomics, *C*. *difficile*

## Abstract

**Background and Aims:** Intratumoral heterogeneity with respect to both host and microbial components has emerged as an important contributor to colorectal cancer (CRC) biology. Although the overall taxonomy and abundance of the CRC tumor microbiome have become well characterized, much less is known about bacterial niches within the tumor microenvironment (TME). *Escherichia coli* and *Fusobacterium nucleatum* are highly prevalent and abundant species linked to CRC. Less abundant organisms, like *Clostridioides difficile* and *Enterocloster aldenensis*, are emerging as potentially important CRC-associated bacteria. Additionally, many studies are confounded by preoperative oral antibiotics or the inclusion of late-stage cancers, both of which may alter the gut microbiome. To better understand the spatial relationships between bacteria and CRC, we generated a molecular atlas based on surgically resected tissue specimens collected from a unique cohort of early-stage CRC patients in whom oral antibiotics were not administered preoperatively.

**Methods:** From 20 CRC specimens, we performed matched histopathological analysis, fluorescence *in situ* hybridization (FISH), whole exome sequencing (WES), 16S rRNA amplicon bacterial DNA sequencing, codetection by indexing (CODEX) multiplex immunofluorescence, and spatial transcriptomics of 756 regions among the specimens. We used 16S rRNA amplicon sequencing data to design and experimentally validate custom bacterial probes that were applied to the spatial transcriptomics. Human colonic organoids were used to validate the relationship between *Clostridioides difficile* toxin B (TcdB) and an E-twenty-six family transcription factor, ELF3.

**Results:** We identified eight intratumoral bacterial niches consisting of a variety of bacterial species, and each niche was associated with unique tumor features. We further defined tumor gene expression patterns correlating with individual bacterial species and show that *C. difficile* has a disproportionate impact on the tumor transcriptome given its relatively low abundance. Specifically, TcdB induces nuclear localization of ELF3, increased cytosolic β-catenin protein, and upregulated WNT signaling.

**Conclusion:** Our study integrates multi-omics to identify bacterial species with biological and spatial relevance in CRC regardless of abundance. These findings will enable further studies to define diagnostic and therapeutic targets for bacteria-associated CRC.

## Introduction

Despite improving therapies, colorectal cancer (CRC) continues to be a leading cause of death worldwide^1^. Evidence suggests toxigenic bacteria may be novel drivers of CRC^2^. For example, *Fusobacterium nucleatum*, *Escherichia coli*, *Bacteroides fragilis*, and *Clostridioides difficile* are associated with CRC in human cohorts and accelerate colonic neoplasia in both susceptible murine and *ex vivo* human models^3–8^. Adjuvant cancer treatments targeting the gut microbiome are under investigation^9,10^, but these efforts lack the specificity needed for precision oncology.

Fecal microbiome studies have associated specific bacterial strains, genes, and metabolites with CRC, but far less is known about the spatial relationship between these microbial species either alone or in biofilms (BF) within the tumor microenvironment (TME)^11,12^. Understanding the relationships between intratumoral bacteria and the progression of CRC is critical for the development of better prediction models and therapeutic strategies.

Multi-omics approaches to mapping CRC have generated novel insights regarding tumor evolution, immune regulation, and metastasis^13–16^. To date, few atlases have characterized intratumoral bacteria as a component of the TME^17^. The relationships between tumor-infiltrating bacteria and CRC biology have just begun to emerge from improved technologies with spatial resolution^4,18–20^.

We generated a multi-omic tumor atlas with a focus on bacterial niches within CRC. We characterized 20 early-stage, treatment-naïve, surgically resected CRC specimens within a larger Malaysian cohort of 120 participants^21–24^. This cohort is unique because individuals did not receive preoperative oral antibiotics, thus preserving bacterial BF and the tumor microbiome. Preoperative oral antibiotics are now standard-of-care, making our cohort specimens a distinctive resource. While the diets and environmental exposures are potentially different than CRC patients from the United States, we have previously shown that colonic and tumor BF patterns are not different between these groups^22^. Additionally, we focused on early-stage CRC (stage I/II) with both precursor lesions and adenocarcinoma present within the same specimens to enable our atlas to define early events in CRC. What emerges from our atlas is the pronounced effect of low-abundance bacterial species on the tumor transcriptional program. *C. difficile* is emphasized as a leading example because of its ability to target the ELF3 transcription factor to the nucleus and to upregulate cytoplasmic β-catenin with WNT pathway activation combined with recent epidemiological evidence that toxin B (TcdB)-positive *C. difficile* associate with the cohort studied herein and positive clinical testing for *C. difficile* toxin B (TcdB) precedes an increase in CRC risk^25,26^.

## Materials and Methods

### Experimental Design

This study was approved by the University of Malaya Medical Centre (UMMC, Kuala Lumpur, Malaysia) Medical Ethics Committee (Ref No. 1066.38) and the Johns Hopkins Institutional Review Board. All samples were obtained between 2012 - 2015 in accordance with the Health Insurance Portability and Accountability Act (HIPAA). The initial cohort consisted of 120 individuals diagnosed with CRC at UMMC, for whom excess tumor and paired normal tissues from the colon were collected following informed consent. Individuals who had received pre-operative radiation, chemotherapy or had a personal history of CRC or inflammatory bowel disease were excluded. All patients underwent a standard osmotic bowel preparation. Standard preoperative intravenous, but not oral, antibiotics were administered in all surgical cases. Demographic and clinical data for this cohort are shown in Supplementary Table 1. After surgical resection at UMMC, excess tumor tissue was collected and preserved via flash freezing, FFPE, and Carnoy’s fixation comprised of methanol:acetic acid:chloroform (6:3:1).

### Bacteria-Tissue Microarray (Bac-TMA)

In total, 41 unique bacterial strains were included in the Bac-TMA (Supplementary Table 2). This included 25 bacterial targets and 13 negative control bacteria. For each target species, at least one control species of the same genus (or similar) was included. In cases where more than one species from the same genus represented a target of interest (e.g., *F. necrophorum*, *F. nucleatum*, *F. periodonticum*, and *F. varium*), these targets also served as negative controls for each other. Strains were acquired from commercial sources (ATCC) or culture collections from institutions listed in Supplementary Table 3. To create the Bac-TMA, bacteria were streaked onto BHI or BRU agar and grown at 37° C anaerobically or aerobically for at least 24 h. Species identity was confirmed using whole 16S rRNA gene Sanger sequencing from bacterial gDNA.

### 16S rRNA Amplicon Sequencing

16S rRNA amplicon sequencing (16S) data of CRC tumor-resected tissues from the Malaysian CRC cohort was compiled from 3 prior Illumina sequencing batches of the V3-V4 hypervariable regions as previously described^21,27^.

### Co-detection by Indexing (CODEX)

Commercially available monoclonal or polyclonal anti-human antibodies were verified by immunofluorescent staining in multiple channels as described previously^28^. Detailed information on antibodies and microscopy is provided in Supplementary Table 4. Antibodies were conjugated with barcodes using the Akoya Antibody Conjugation kit (SKU 7000009). CODEX staining and imaging were performed according to the manufacturer’s instructions (CODEX user manual, Rev C).

### Spatial Transcriptomics

Tissue blocks were sectioned at 5 μm thickness onto standard slides under RNase-free conditions using RNase AWAY (ThermoFisher #7002). The same preparation protocol was used for both the tumor tissue and the Bac-TMA (GeoMx DSP Manual Slide Preparation, MAN-10150-04) as we performed previously^29^. Pan-cytokeratin, CD45, and SYTO13 morphology markers were used to identify tissue features. The human Whole Transcriptome Atlas (WTA) probe set was used to hybridize host tumor transcripts, and UV-cleaved oligonucleotide barcodes were sequenced to generate gene counts.

### Colonic Organoids (Colonoids)

Normal human colonoids were derived from biopsies taken during routine colonoscopies of healthy adults (24 - 72 years old) at the Nashville VA Medical Center (IRB #1523236). All individuals provided informed consent for biopsies of normal tissue to be used for research purposes, specifically colonoid generation. Colonoids were maintained at low passage numbers embedded in Matrigel (Corning) domes submerged in Intesticult Organoid Growth Media (StemCell Technologies) as described previously^29^. Experiments were repeated in colonoids from two separate individuals (biological replicates) using independently passaged domes (technical replicates) as indicated in each experiment.

### Statistical Analysis

The Kruskal-Wallis test, Dunn’s test of multiple comparisons, χ^2^ testing, and standardized residuals were prepared using the *rstatix* (v0.7.2) package in R (v4.4.2). Spearman correlations were generated using the *stats* (v4.4.2) package. Power calculations were performed using the *pwrss* (v0.3.1) package. Unless otherwise stated, plotting was performed using the *ggplot2* (v3.5.1) and *ggpubr* (v0.6.0) packages.

See the Supplementary Materials for more details on FISH and biofilm screening, the Bac-TMA, 16S, qRT-PCR, CODEX, spatial transcriptomics analysis, and image registration.

## Results

To build on our HTAN atlas mapping precancer-to-cancer transitions^13^, we examined early-stage CRCs that captured precursor lesions and adenocarcinoma within the same specimens. We examined 3 groups: BF^+^ right-sided, BF^+^ left-sided, and BF^-^ left-sided (Figure 1A). Nearly all right-sided tumors in the Malaysia cohort are BF^+^ (>90%)^21^. The participants were 55% women (n = 11), ranged in age from 52 to 84 years, and self-identified as Chinese (65%), Malay (20%), and Indian (15%) (Supplementary Table 1). Portions of each CRC specimen were preserved separately via flash freezing, Carnoy’s fixation, and formalin-fixation, paraffin-embedding (FFPE) (Figure 1B). We identified invasive BF and intratumoral bacterial colonies using multiprobe-fluorescence *in situ* hybridization (FISH) on Carnoy’s fixed tissue and FFPE samples. Hematoxylin and eosin (H&E), codetection by indexing (CODEX), and GeoMx spatial transcriptomics were performed on serial sections of FFPE specimens.

**Figure 1.**
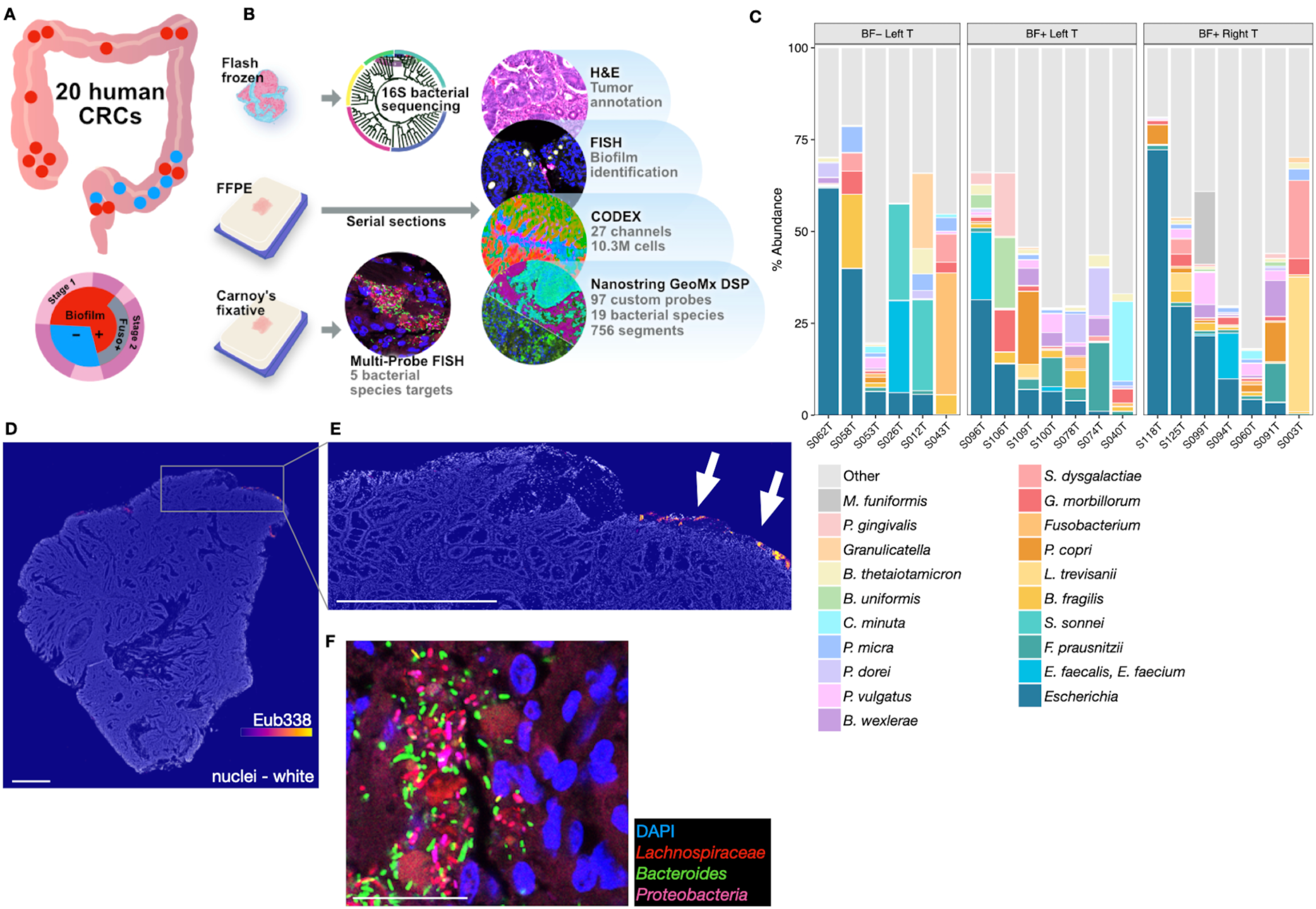
Tumor surface biofilms and bulk 16S sequencing. (*A*) Overview of sampling and workflow for the nested cohort comprising 20 tumors with matching multi-omics. This study encompassed 10 tumors from the right colon (red dots are biofilm positive [BF^+^]) and 10 from the left colon. The circle diagram represents the proportions of the cohort with tumor characteristics: BF-status, presence of *Fusobacterium* blooms, and tumor stage. (*B*) Cryopreserved specimens were used for bulk 16S sequencing. Carnoy’s fixed samples were used for multi-probe FISH and confocal microscopy. FFPE specimens were cut for FISH, CODEX, H&E, and GeoMx with bacterial species-specific probes. (*C*) 16S data reveal the most abundant species in each bulk flash-frozen tumor specimen. (*D*) Stitched microscopy image of tumor S109 stained with fluorescence in situ hybridization (FISH) and pseudo-colored by the intensity of a universal 16S probe (Eub338); scale bar = 1 mm. (*E*) Inset magnification of the white-boxed area from (*D*); scale bar = 1 mm. (*F*) Inset magnification of multi-probe FISH microscopy image from Supplementary Figure 1C with polymicrobial biofilm; Red = *Lachnospiraceae*, Green = *Bacteroides*, Pink = *Proteobacteria*, Blue = nuclei.

We analyzed the 16S results and found no clear clustering of species-level microbiome composition by BF status or tumor location (Figure 1C, Supplementary Figure 1A). These data were consistent with our prior reports in larger cohorts showing BF status is not readily predicted by sequencing alone^21,22,27^. At the species level, half (10/20) of the tumors had a relatively high abundance (> 10%) of *E. coli*, *E. faecalis*, or *Fusobacterium* spp. Next, we examined the BF status of each tumor from both Carnoy’s fixed and formalin-fixed specimens and observed discontinuous BF localization on the tumor surfaces (Figure 1D-E). Previously, we identified four predominate taxonomic groups in CRC-associated biofilms: the Bacteroidetes phylum (including *Bacteroides, Parabacteroides,* and *Prevotella*)*, Fusobacterium* spp., Lachnospiraceae family, and Gammaproteobacteria and Betaproteobacteria classes^21^. In the present cohort, we examined these groups using multi-probe FISH on Carnoy’s fixed tissue. All BF^+^ tumors had polymicrobial biofilms (Figure 1F, Supplementary Figure 1B-C). Half of the BF^+^ tumors (7/14) contained *Fusobacterium* genus with a range from dense blooms in 5/14 BF^+^ tumors to fewer than 5 bacteria in a 0.04 mm^2^ field of view. Given the variability in density and location of bacteria within these heterogeneous tumors, our data suggest spatial resolution is required to fully understand the relationship between bacteria and CRC biology.

To incorporate bacterial abundance for this spatial CRC atlas, 16S results were used to generate a list of target species. We designed three custom bacterial probes for each of 19 target species that were either highly abundant or found in previously identified CRC-associated BFs^8^. In addition, we designed eight “universal” probes to hybridize with conserved regions of the 16S rRNA. We also included 32 negative probes designed without any known bacterial or human target to generate a background noise threshold. Similarly to Zhu et al^30^, we validated these “spike-in” probes on the GeoMx platform with a bacteria-tissue microarray (BacTMA) containing cultured isolates of sequence-verified bacteria and colonic tissue from gnotobiotic mice (Supplementary Figure 2A). We applied the BacTMA to the GeoMx platform with the custom probes to determine the appropriate sensitivity and specificity for tumor analysis (Supplementary Figure 2B, Supplementary Table 2). Heatmaps for each bacterial probe set were generated to analyze the detection of targets within the BacTMA. For example, all three probes against *Phocaeicola* (*Bacteroides*) *dorei* were sensitive for detection, but only *P. dorei* probes 02 and 03 demonstrated acceptable specificity for inclusion in the tumor experiments (Supplementary Figure 2C-E). This custom, validated set of species-specific probes was applied to investigate microbe-host interactions in human CRC tumor tissue.

### Spatial niches of bacteria in CRC

To identify spatial regions of interest (ROIs), we first examined H&E- and FISH-stained FFPE serial sections for histopathological features and bacteria-rich zones, respectively. Within each tumor, histological features were designated by expert gastrointestinal pathologists as the following: normal, adenomatous precursor lesions, foci of high-grade dysplasia (HGD), intramucosal adenocarcinoma (AC), and invasive AC. We applied both the validated bacterial probes and the GeoMx human WTA to serial FFPE sections. We sought to balance the ROIs based on FISH score (semi-quantitative measurement of total bacteria in a region), intratumoral location, and epithelial versus non-epithelial cell types (Figure 2A, Supplementary Figure 3A). We focused attention on ROIs in transition zones that consisted of juxtaposed normal, precursor, and/or adenocarcinoma) (Figure 2B). Each GeoMx ROI was further segmented based on pancytokeratin immunofluorescent staining into epithelium (PanCK^+^) and non-epithelium (PanCK^-^) tissue. All together, we collected 756 ROIs from the 20 tumors. The H&E and FISH images were then registered with GeoMx and CODEX images to demonstrate ROIs that targeted overlapping areas of bacteria and tumor pathology; for example, a high bacterial load near an ulcerated tumor surface is shown from sample S078 (Supplementary Figure 3B).

**Figure 2.**
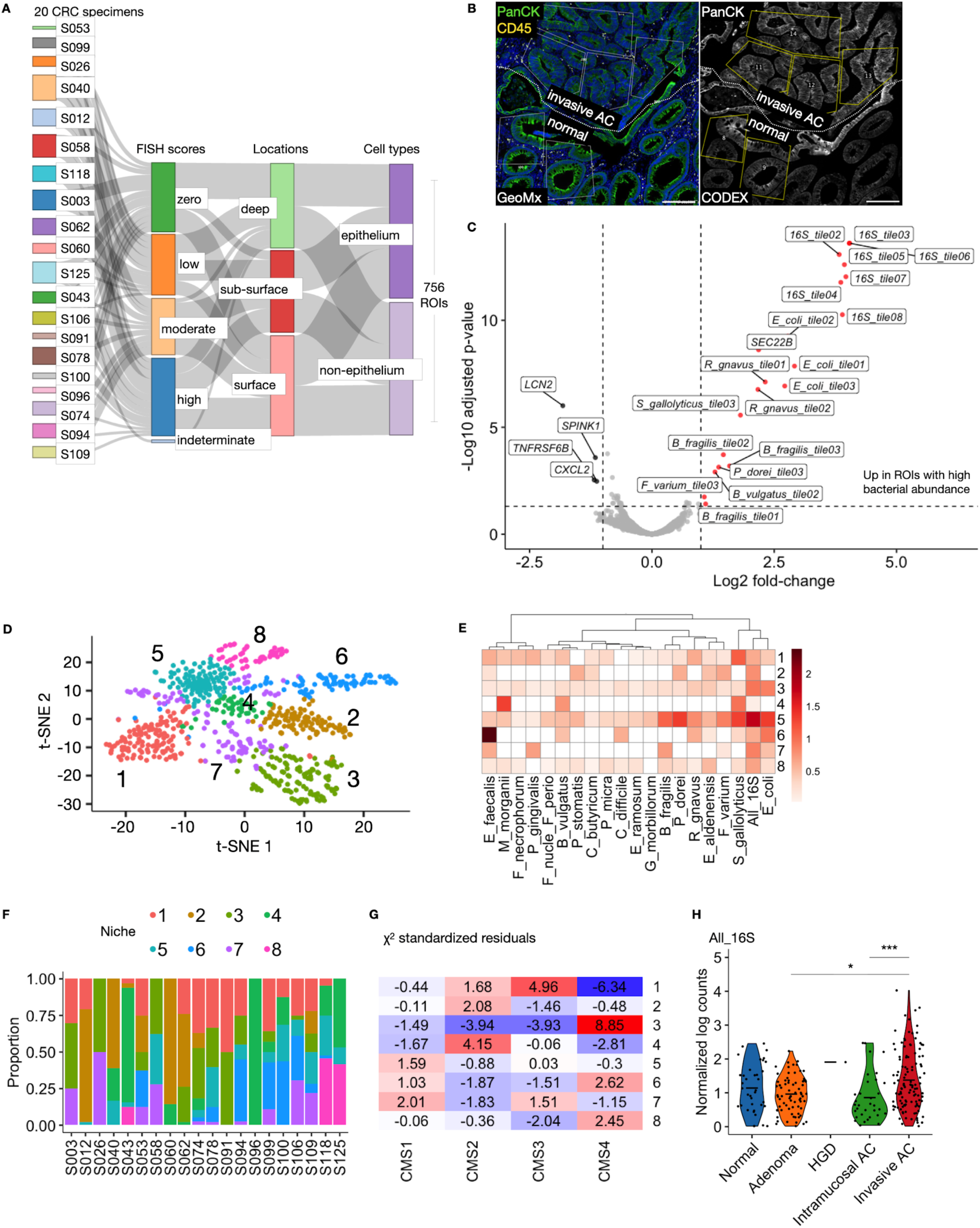
Characterization of bacterial niches within CRC. (*A*) Sankey diagram illustrating the characteristics of all 756 regions of interest (ROIs) chosen for spatial transcriptomics. (*B*) Fluorescence microscopy from serial sections demonstrating the high fidelity of image registration and mapping of GeoMx spatial transcriptomics ROIs onto CODEX images. The dotted line marks a transition zone between normal-appearing crypts and invasive adenocarcinoma (AC). PanCK = pancytokeratin. (*C*) Volcano plot comparing the differential expression or abundance of genes and bacterial probes in ROIs with high bacterial load (right side) to ROIs with low bacterial load (left side). (*D*) t-distributed Stochastic Neighbor Embedding (t-SNE) plot of the GeoMx ROIs informed by both the WTA and the custom bacterial probes. Louvain clustering identified eight different niches. (*E*) Heatmap with hierarchical clustering that shows the normalized, log-transformed median abundance for each species detected by the custom bacterial probes in each niche. (*F*) Relative proportions of niches by tumor. (*G*) Bacterial niches and CMS classification for tumors are significantly correlated (χ^2^= 148.43, p-value = 0.0005). χ^2^ standardized residuals demonstrating specific niches are significantly enriched (residuals > 1.96, p < 0.05, highlighted in red) or depleted (residuals < - 1.96, p < 0.05, highlighted in blue) when correlated with CMS classifications. (*I*) Violin plot comparing the abundance of total bacteria (geometric mean of eight universal 16S probes normalized to background and log-transformed) across different histopathological features within the CRC tumors; p-values obtained from Dunn’s test: \**p* < 0.05, \*\*\**p* < 0.001.

To define the features associated with high bacterial abundance, we binned the ROIs into the highest and lowest quartiles based on normalized mean counts of total bacteria as detected by the eight universal probes hybridizing with conserved portions of the 16S rRNA (All_16S). We then performed differential expression analysis that revealed only one host gene significantly upregulated in ROIs with high bacterial abundance, *SEC22B* (Figure 2C). The autophagy-associated vesicle adaptor *SEC22B* can promote CRC cell growth under stressful conditions and is associated with MSS CRC risk^31^. The remaining 19 significantly enriched transcripts detected were of microbial origin. As expected, seven were detected by the universal All_16S probes. However, 12 species-specific probes representing 7/19 different species were also significantly enriched in ROIs with the highest total bacterial abundance. These data suggest bacteria-rich ROIs in early-stage cancers are generally heterogeneous. The relative expression of *LCN2* and *CXCL2*, which are associated with infiltrating neutrophils, in ROIs with low bacterial abundance suggests that an active innate immune response may limit bacterial accumulation^32,33^.

Next, we used the WTA and bacterial probe counts to perform dimensionality reduction with t-distributed stochastic neighbor embedding (tSNE) and Louvain clustering to show that the 756 GeoMx ROIs clustered into 8 niches (Figure 2D)^34,35^. We observed more heterogeneity with respect to the presence of each individual targeted species within the niches (Figure 2E). There were some exceptions, like *E. faecalis* enriched in Niche 6 and *M. morganii* abundant in Niche 4 (Supplementary Figure 4A). *E. coli* was highly enriched in Niche 5, which also had the highest mean abundance of *S. gallolyticus*, *P. dorei*, and *B. fragilis* (Figure 2E). Furthermore, Niche 5 had the most bacteria detected by the All_16S probes and was the only niche containing each of the 19 species we targeted (Supplementary Figure 4A-B). Niche 4 had the fewest number of different species. Overall, each tumor harbored a heterogeneous set of microbial niches (Figure 2F).

Notably, GeoMx ROIs did not segregate well by other factors, including tissue type, FISH score, or location within a given tumor (Supplementary Figure 4C-I). However, there was some clustering based on epithelium versus non-epithelium and tumor (Supplementary Figure 4E, 4I). Further analysis by hierarchical clustering suggested the ROIs clustered modestly by epithelium versus non-epithelium (Supplementary Figure 4J). Expectedly, cell type appears to be a main driver of the ROI clustering.

Next, we used the GeoMx data to perform consensus molecular subtyping (CMS) predictions for each ROI (Supplementary Figure 5A). Even though 18/20 tumors exhibited heterogeneity in CMS prediction (Supplementary Figure 5B), *E. coli* significantly correlated with CMS3 (positive) and CMS1 (negative) (Supplementary Figure 5C). Of the 8 niches, there were significant associations with particular CMS classifications (Figure 2G), including the high degree of overlap between Niche 1 to CMS 3, Niche 3 to CMS4, and Niche 4 to CMS2. Niche 6 associated with CMS4, an aggressive subtype, and was enriched with *E. faecalis* (Figures 2E, 2G). This association is consistent with other data linking *E. faecalis* production of a collagenase enzyme through expression of the *gelE* gene with cancer cell invasion^36^.

Next, we sought to characterize further the TME of ROIs with the highest bacterial abundance. With respect to histopathological features, the areas of invasive AC were generally most enriched with bacteria (Figure 2H). The ROIs within or bordering tumor necrosis also contained a high bacterial load (Supplementary Figure 6A-C), which has been previously reported^17^. *F. varium*, *P. dorei*, *R. gnavus*, and *S. gallolyticus* are particularly abundant in necrotic ROIs compared to non-necrotic ROIs (Supplementary Figure 6B). We compared the differential gene expression from the epithelium bordering necrotic areas to all non-necrotic areas in the dataset (Supplementary Figure 6D), and we observed increased expression of *CXCL8*, which has been previously shown to be upregulated within bacteria-rich ROIs in CRC^17,20^. The relatively lower bacterial load in non-necrotic areas was associated with the expression of immunoglobulin-producing genes *IGKC* and *IGHG3*. Gene set enrichment analysis (GSEA) comparing necrotic to non-necrotic areas revealed a relative lack of response to pathogens in the necrotic areas (Supplementary Figure 6E). For example, the tumor necrosis factor (TNFα) and interferon-γ responses were enriched more in non-necrotic areas, concordant with fewer bacteria. Interestingly, the enriched transcriptional programs in necrotic areas were inferred to activate epithelial-to-mesenchymal transition, consistent with cancer progression and prior reports that define necrosis as an active process involving neutrophil-driven vascular occlusion (Supplementary Figure 6E)^37^.

The individual probes within the probe sets were compared by hierarchical clustering, which showed probes targeting the same species clustered closely together (Supplementary Figure 7A). This analysis supported our validation of the probes and provided confidence for using the mean of each probe set. Generally, only positive correlations were detected between bacterial abundance across all the custom bacterial probe sets (Supplementary Figure 7B). We also tested for correlations between the detection of these probe sets from GeoMx and the bulk tumor 16S data from bulk frozen portions of the same tumors (Supplementary Figure 7C). Notably, co-detection by 16S and GeoMx was statistically significant for *B. fragilis*, *P. gingivalis*, *P. micra*, and *S. gallolyticus*, despite these data originating from two different segments of each tumor.

We also performed targeted qPCR for key procarcinogenic microbes. All tumors were positive for *E. coli*, and 12/20 (60%) tumors were positive for *B. fragilis* 16S (Supplementary Figure 8A-B). Then, we tested more specifically for toxin-producing genes and found 7/20 (39%) tumors were positive for *pks^+^ E. coli*, and no tumors had detectable *bft^+^ B. fragilis* (Supplementary Figure 8C-D). Others have demonstrated *bft^-^ B. fragilis* and *pks^-^ E. coli* may have alternative mechanisms influencing carcinogenicity^38^. We previously showed in a meta-analysis of 16S datasets that *B. fragilis* as a species (regardless of *bft* status) was significantly enriched in CRC when compared to paired normal or healthy controls from assessment of both stool and colon tissue in 13 international cohorts^21^. We did not see differences in *E. coli* from that 16S meta-analysis. Similarly, Wirbel et al. did not report enrichment of *E. coli* in their large fecal metagenomics CRC study^39^. Furthermore, the reported prevalence of the *pks* island in CRC is highly variable (10% - 55%)^3,40,41^. The prevalence of *bft* in CRC cohorts also varies widely from 27% to 85%^42,43^. In one published cohort, *bft* was not significantly enriched in CRC versus paired normal tissue^39^. These results suggest *pks^-^ E. coli* and non-enterotoxigenic *B. fragilis* are common in this cohort, and our current understanding of variability in detection of bacterial virulence factors in CRC remains incomplete.

### Specific bacteria in the TME

*E. coli* was the most abundant species in the cohort, and the Gram-positive species *R. gnavus* and *S. gallolyticus* were highly represented across multiple histopathological tissue types (Figure 3A-B). Interestingly, HGD was enriched with bacteria but not any of the 19 specifically targeted species (Figure 3B). This result may be skewed by the relatively few ROIs in areas of HGD. Most tumors were heterogeneous, while others were dominated by a single species, like S096 with *S. gallolyticus* and S118 with *E. faecalis* (Figure 3C). The custom bacterial probes were able to detect species in ROIs that scored “Zero” by FISH (Figure 3D), demonstrating the high sensitivity of our custom probes. Surprisingly, there was higher general abundance of species present deep within tumors compared to the sub-surface ROIs, consistent with prior data showing extensive tumor invasion by bacteria in BF^+^ CRC (Figure 3E)^21^. Importantly, our FISH methodology did not distinguish between intracellular and extracellular bacteria. Some species, like *E. coli*, appeared to be evenly distributed between the epithelium and non-epithelium, but several other species (e.g. *S. gallolyticus*, *B. fragilis*, *F. varium*, *B. vulgatus*, and *R. gnavus*) were more highly associated with the non-epithelial segments (Figure 3F), which could reflect infection or phagocytosis of the microbes by immune cells.

**Figure 3.**
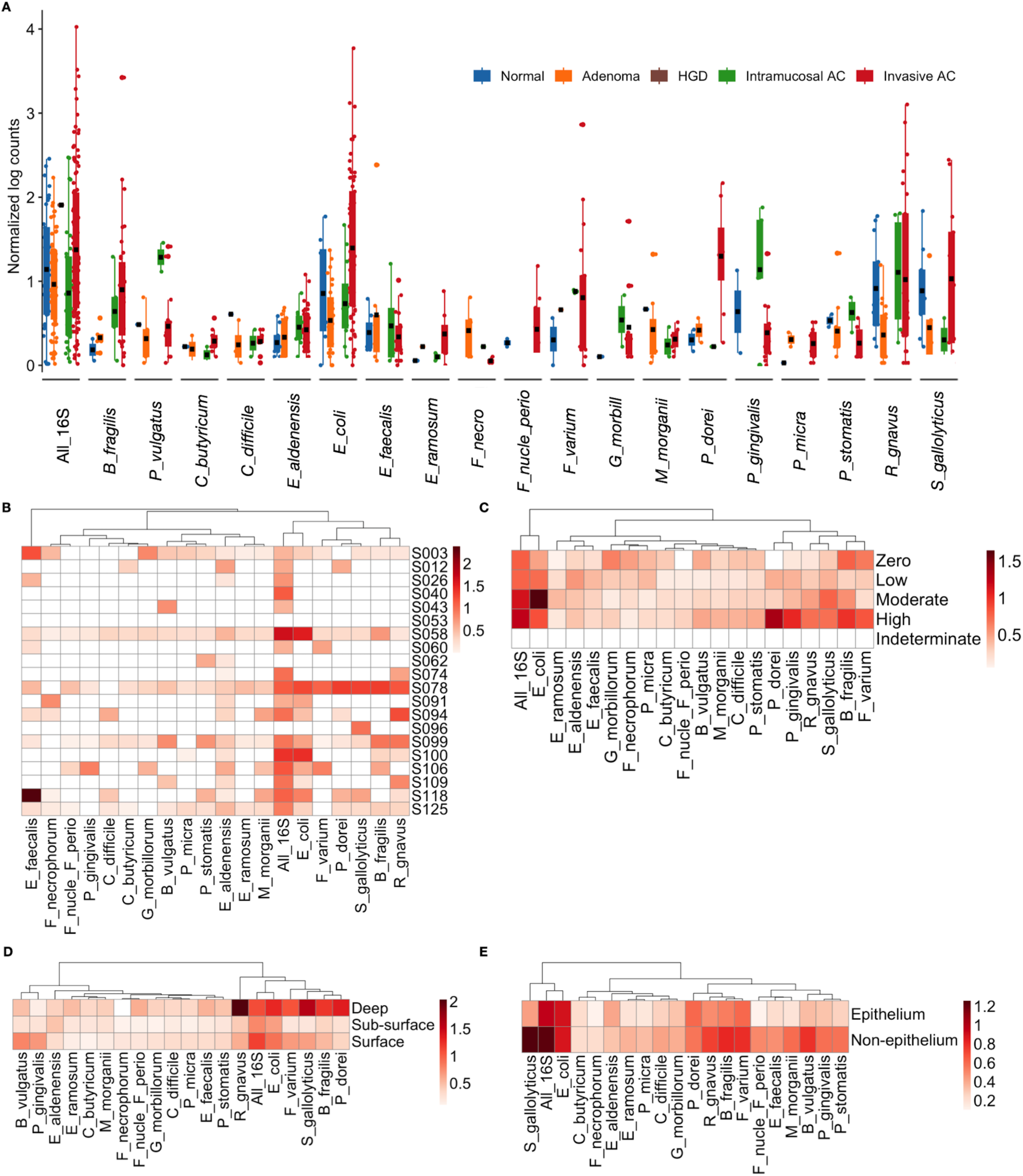
Bacterial species abundance within tumor features. (*A*) Bar-whisker plot of bacterial species abundance colored by the histopathological tissue types; adenocarcinoma (AC). Black dots represent the mean; whiskers are top and bottom quartiles. (*B*) Heatmap with the median normalized, log-transformed probe count for each custom bacterial probe set. (*C*) Median bacterial species count by tumor. (*D*) Median bacterial species count by FISH score (see Supplementary Figure 3A). (*E*) Median bacterial species count by location within tumors. (*F*) Median bacterial species count by epithelial versus non-epithelial segments.

### Tumor transcriptional programming in bacterial niches

Moving from the tissue level to classification of the bacterial niches at the molecular level, we performed differential gene expression analysis for each niche compared with all the other niches. More specifically, we focused on the PanCK^+^ epithelial segments from dysplastic regions of the tumors. Niche 1 was enriched with immunoglobulin genes, such as *IGKC*, *IGHG1-4*, and *JCHAIN*, suggesting a relatively high infiltration of plasma cells into the epithelium (Supplementary Figure 9A). The *MZT2B* gene was upregulated in Niche 2 but downregulated in Niche 3 (Supplementary Figure 9B-C). Although there are no reports related to CRC, *MZT2B* is a MYC-regulated gene previously demonstrated as a marker of aggressive intestinal-type gastric cancer^44^. Niche 3 exhibited upregulation of ribosomal genes, while Niche 4 showed relative downregulation of ribosomal genes (Supplemental Figure 9C-D). The *IGF2* gene was significantly upregulated in Niches 5 and 8, while Niche 6 exhibited bacterial probe abundance and *SEC22B* upregulation similar to the ROIs with high total bacterial load (Supplementary Figure 9E-H, Figure 2C). *IGF2* is a mitogen associated with an increased risk of developing colonic neoplasia^45^. *NXPE4* is among the few genes significantly upregulated in Niche 7, and it has previously been suggested as a prognostic biomarker for CRC and may be capable of sialoglycan acetylation in the colon (Supplementary Figure 9G)^46,47^. We used GSEA from both the Hallmark MSigDB and gene ontology gene sets to identify global trends for niche functionality (Figure 4A, Supplementary Figures 10-11)^48^. Niches 1, 3, and 7 are the most enriched with the Hallmark signaling pathways, although none of the normalized enrichment scores were statistically significant for Niche 7; the remaining Niches (2, 4, 5, 6, and 8) were de-enriched for these pathways (Figure 4A). One explanation for this demarcation between the two groups of niches could be that the ROIs comprising Niches 1, 3, and 7 were more likely to be intramucosal and invasive AC (Figure 4B). The relatively few associations with HGD are potentially explained by the paucity of HGD tissue in these tumor specimens.

**Figure 4.**
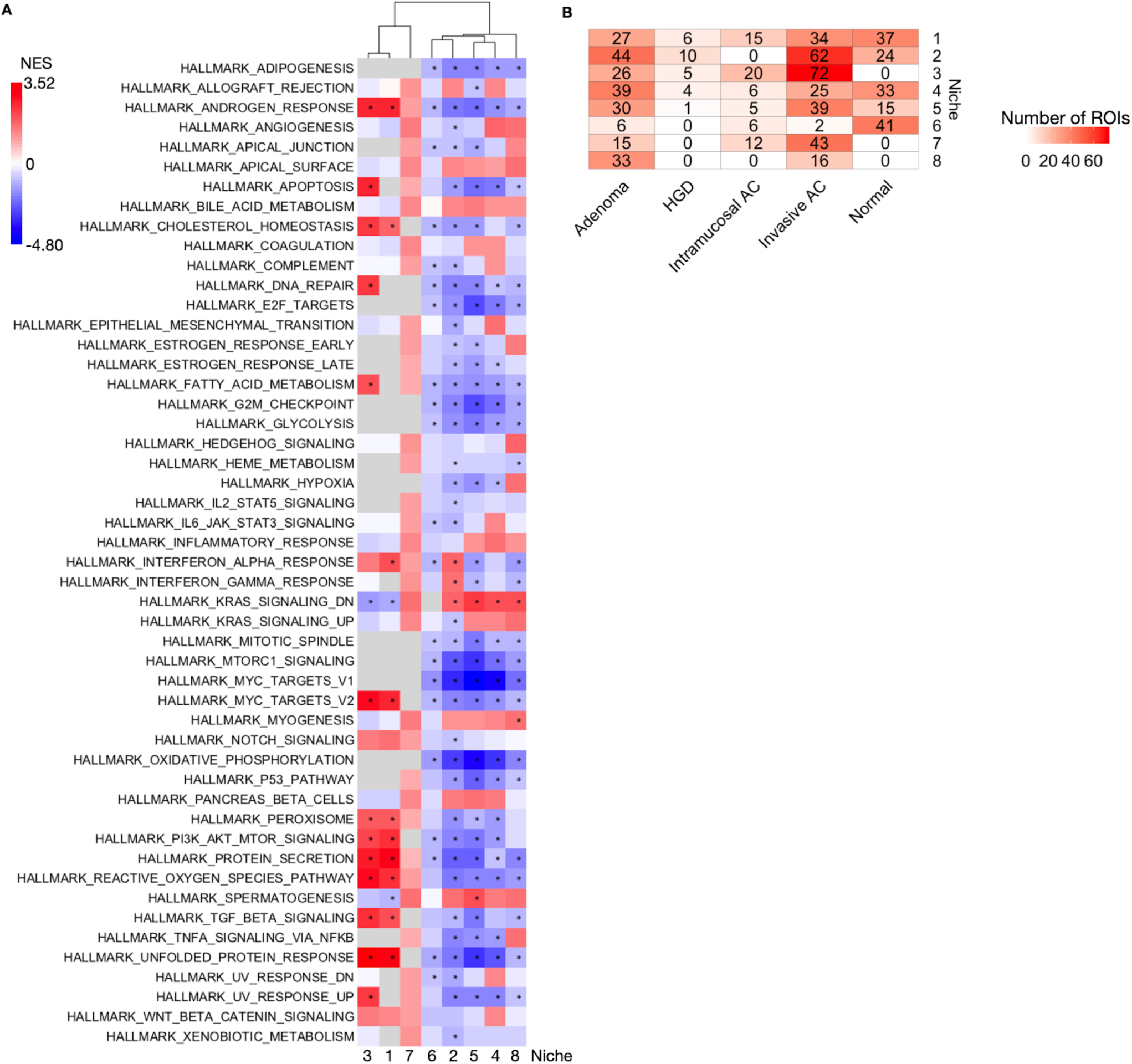
Gene set enrichment analysis (GSEA) characterization of niches. (*A*) Heatmap showing the enrichment of Hallmark pathways from the Human Molecular Signatures Database (MSigDB). Normalized Enrichment Score (NES) as defined previously^63^. Gene sets for GSEA were based on the differential gene expression analysis illustrated in Supplementary Figure 9. (*B*) Heatmap of the raw number of ROIs from each niche that are within the histopathologically defined tumor features.

### Immune infiltration and bacterial niche localization

To characterize bacterial niches at the cellular level, we analyzed CODEX images of entire tumor tissue serial sections with a focus on immune cell infiltration within similar ROIs as used for GeoMx (Figure 5A-B, Supplementary Figure 12A). Cell segmentation was based on membrane markers, and UMAP dimensionality reduction and Leiden clustering identified cell type assignments in combination with marker expression patterns (Figure 5C, Supplementary Figure 12B). Cell types were enumerated for each bacterial niche (Figure 5D-E). To validate the CODEX data, we applied spatial deconvolution to the GeoMx dataset. These data corresponded with our CODEX cell counts (Supplementary Figure 13A), particularly in reflecting the distribution of Tregs and CD8^+^ T cells in the niches. Next, we evaluated the correlation between specific cell types from CODEX data with specific bacterial abundance from the GeoMx data (Figure 5F-G). Most species had a statistically significant positive correlation with epithelial cells (PanCK^+^), but not proliferating epithelial cells as determined by Ki67 staining. The probes with the highest epithelial cell correlation coefficients were All_16S, *E. coli*, *E. aldenensis*, and *C. difficile* (Figure 5F). For the non-epithelial cell populations, we observed a slight but significant negative correlation between All_16S and CD8^+^ T cells (Figure 5G). In contrast, macrophages correlated positively with All_16S, *E. coli*, *M. morganii*, and *C. difficile*. *M. morganii* was also associated with CD4+ T cells. *P. gingivalis*, *E. faecalis*, and *E. aldenensis* were all associated with mixed T cells (Figure 5G). We were surprised to find significant correlations between low-abundance bacteria, like *C. difficile* and *E. aldenensis*, and specific host tumor cell types, which we investigated further in the next section.

**Figure 5.**
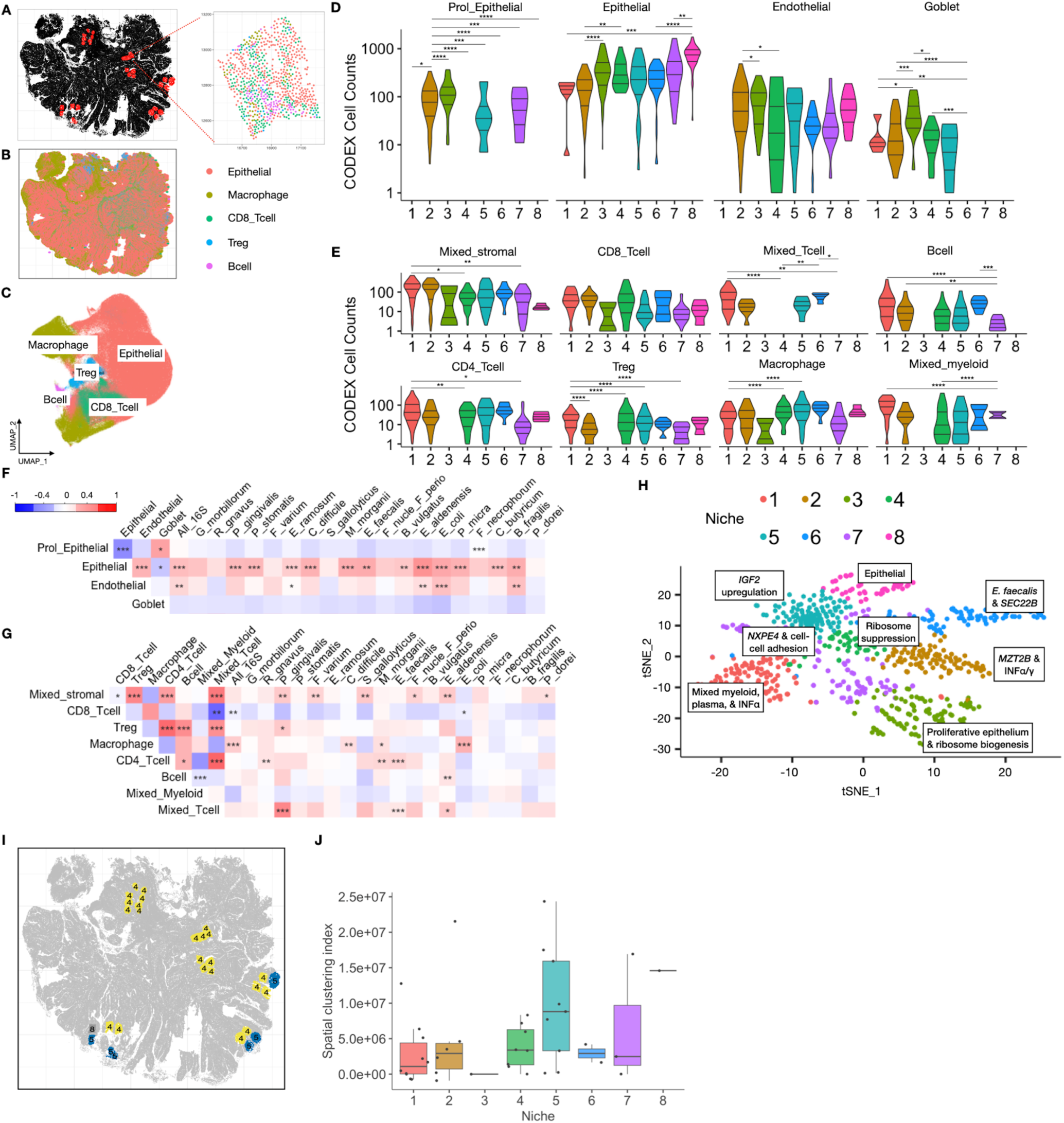
Tumor cell typing and spatial clustering of bacterial niches. (*A*) Spatial map of tumor S125 with GeoMx ROIs in red and one example magnified inset colored by specific cell types derived from co-detection by indexing (CODEX) multiplex immunofluorescence. (*B*) Map of cell types throughout the whole S125 tumor. (*C*) Uniform Manifold Approximation and Projection (UMAP) plot of the S125 tumor cell types based on the expression of all 27 markers (Supplementary Figure 12B, Supplementary Table 4). (*D*) Violin plots of the epithelial cell counts for each niche. \**p* < 0.05, \*\**p* < 0.01, \*\*\**p* < 0.001, \*\*\*\**p* < 0.0001. (*E*) Violin plots of the stromal and immune cell counts for each niche. (*F*) Spearman correlation plots demonstrating the association of bacterial species abundance from GeoMx with epithelial cell type counts from CODEX; \**p* < 0.05, \*\**p* < 0.01, \*\*\**p* < 0.001. (*G*) Spearman correlation plots for the association of bacterial species and non-epithelial cell types; \**p* < 0.05, \*\**p* < 0.01, \*\*\**p* < 0.001. (*H*) Labeled UMAP plot with simplified niche characterizations based on unique features from GeoMx and CODEX data. (*I*) S125 tumor map with spatial distribution of GeoMx ROIs labeled by niche demonstrating the self-colocalization of niches. (*J*) Box-whisker plot of spatial clustering of niches evaluated by Ripley’s K indices, previously shown to be well-powered to detect spatial co-clustering patterns^64^. The indices were normalized for random “null” patterns; the observed positive values for all niches thus indicate that self-clustering is greater than expected and not randomly distributed. These results were not sensitive to the choice of co-clustering radius (500, 1000, or 2000 pixels; 1000 is shown for the figure) in Ripley’s K calculations.

After integrating the GeoMx and CODEX datasets to correlate cell types and bacterial abundance, we were able to identify unique traits associated with each niche (Figure 5H, Supplementary Table 5). These niche assignments are specific but not fully comprehensive. Niche 1 was identified as being enriched with mixed myeloid and plasma cells with upregulation of the interferon-α (INFα) signaling pathway. Significant upregulation of both interferon-γ and INFα (Figure 4A), along with the *MZT2B* gene (Supplementary Figure 9B), were present in Niche 2. Niche 3 showed remarkable enrichment of the ribosomal biogenesis pathways from GSEA (Supplementary Figure 10C) and proliferative epithelium from the CODEX analysis (Figure 5D). Conversely, suppression of ribosome complexes was predicted for Niche 4 (Supplementary Figure 10D). Niche 5 notably contained a single upregulated gene, *IGF2*, from the differential gene expression analysis of the epithelial segments in dysplastic areas (Supplementary Figure 9E). The upregulation of *SEC22B* and the high abundance of *E. faecalis* uniquely marked Niche 6 (Supplementary Figure 9F, Figure 2E). Niche 7 showed enrichment of genes related to cell-cell adhesion and upregulation of the CRC biomarker *NXPE4* (Supplementary Figure 9G, Supplementary Figure 11C). The high number of total epithelial cells was the predominant factor for Niche 8 (Figure 5D), which also was made up of mostly adenoma and invasive AC cells (Supplementary Figure 4C). These unique niche characteristics will be interrogated with more mechanistic hypotheses in future studies.

We next mapped all the spatial location of the niches onto the tumor and calculated a spatial clustering index (Ripley’s K) to determine if niches co-localize with themselves or localize preferentially with other niches^49^. Niches of the same type tended to co-localize (Figure 5I-J), and different niche pairs were co-excluded from one another (Supplementary Figure 13B). For example, ROIs from Niche 1 co-localize with other Niche 1 ROIs but are separated from ROIs of Niche 2. These niches were derived from GeoMx ROIs chosen to represent histopathological transition zones. Therefore, it appears the bacterial abundances and transcriptional programs are similar among niches that are close together despite spanning different histopathological features.

### Bacterial abundance and tumor gene expression

Next, we correlated species detection with specific gene expression. *E. coli* was an expected species to stand out in these results because of its overall high abundance in our cohort (Figure 6A). Intriguingly, the highest correlative expression with *E. coli* abundance was the previously mentioned autophagy-associated, vesicle adaptor *SEC22B*. Surprisingly, abundance of *E. aldenensis* and *B. fragilis* was associated with the expression of many tumor genes (Figure 6A, Supplementary Table 6). These species were not among the most highly abundant species in our cohort, suggesting their biological relationship with the tumors was stronger than predicted by abundance alone. In fact, other species with low overall abundance were significantly correlated with tumor gene expression: *C. difficile*, *M. morganii*, and *P. micra* (Figure 6A). To examine these relationships more carefully, we plotted the significant positive and negative correlating genes on a volcano plot for All_16S and each species separately (Figure 6B, Supplementary Figure 14, Supplementary Table 6). Among the genes with the highest absolute correlation coefficient was *SEC22B*, and we illustrate its relationship with total bacterial abundance in Figure 6C. We next used the web-based package ChIP Enrichment Analysis 3 (ChEA3) to predict a transcription factor regulatory network associated with the top 100 genes positively correlated with the All_16S probes (Figure 6D)^50^. Because of our prior interest in *C. difficile*-associated tumorigenesis and its previously mentioned association with specific cell types, we repeated the same analysis for *C. difficile* abundance and tumor gene expression (Figure 6E-G). There was a slight but significant positive relationship between the expression of *TEX2* and the detection of *C. difficile* (Figure 6F). *TEX2* gene methylation has been implicated in high-risk CRC^51^. Interestingly, the transcription factors ELF3 and FOSL1 appeared for both the All_16S and *C. difficile* regulatory networks (Figure 6D, 6G). Taken together with our prior data in mouse models and human cohorts^25,26^, we hypothesized that *C. difficile* toxin B (TcdB) activates a pro-tumorigenic transcriptional network.

**Figure 6.**
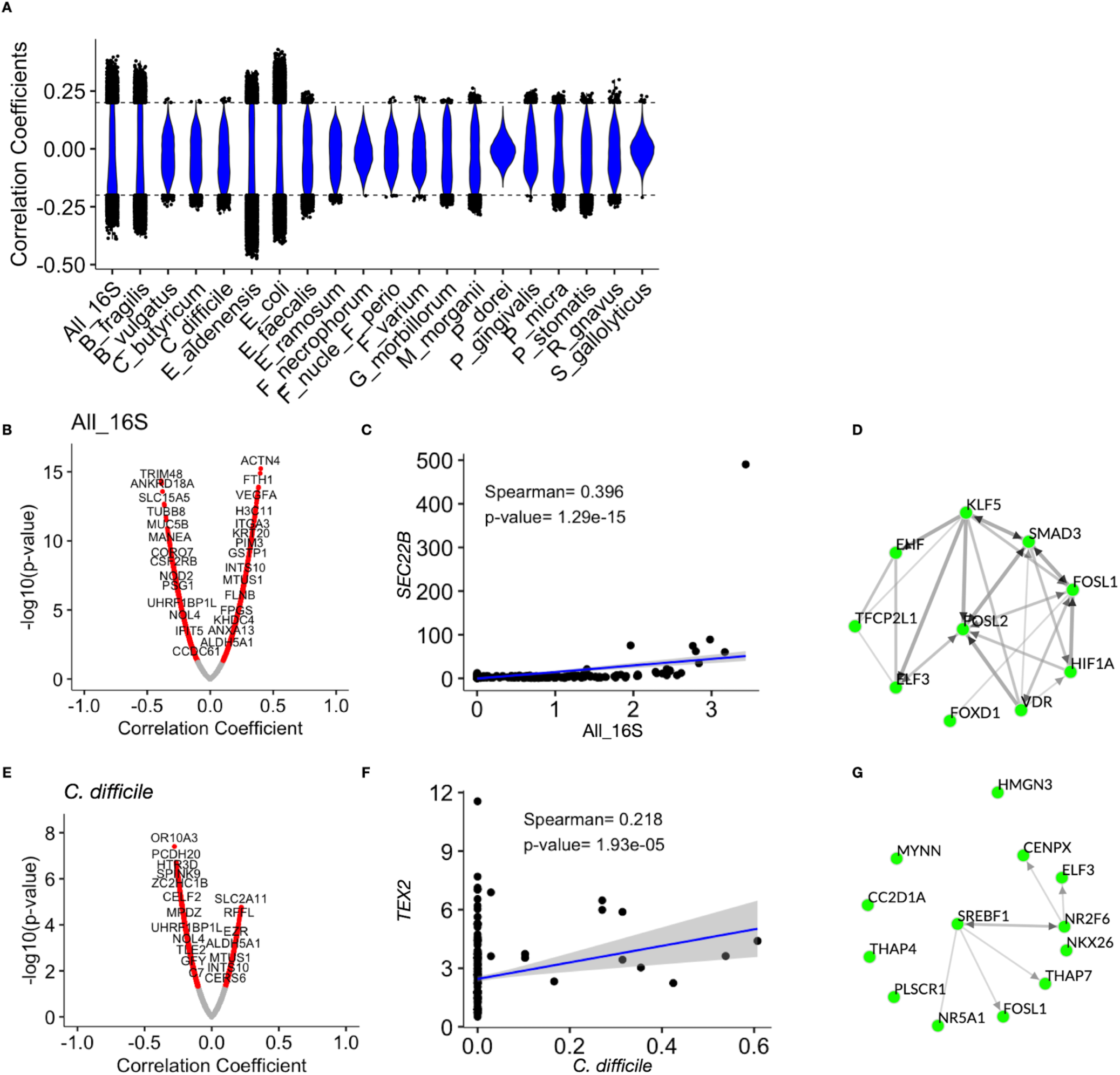
Integration of bacterial species counts with gene expression data. (*A*) Violin plot showing the correlation coefficients between bacterial abundance for each species (separate columns) and the expression of each tumor gene (black dots denote genes with a correlation coefficient greater than 2 or less than -2). (*B*) Volcano plot of the specific genes with statistically significant correlations between their expression and the abundance of total bacteria. (*C*) The correlation between *SEC22B* gene expression and the abundance of total bacteria (normalized, log-transformed) is shown as a scatter plot in which each dot represents a GeoMx ROI. (*D*) Inferred regulatory network of transcription factors for the top 100 positively correlated genes from (*B*) derived using chromatin immunoprecipitation enrichment analysis (ChEA3)^50^. (*E*) Volcano plot for the correlations between *C. difficile* abundance and specific gene expression. (*F*) Scatter plot of GeoMx ROIs demonstrating the correlation between *TEX2* expression and *C. difficile* bacterial counts. (*G*) ChEA3-derived transcription factor regulatory network for genes positively correlated with *C. difficile*.

### Validation of ELF3 transcriptional activation by TcdB

Recently, we published an analysis of *C. difficile* prevalence in the larger Malaysian cohort and showed 38% of individuals had CRC tumors with detectable *C. difficile* with 22% of 65 tissues examined positive for *tcdB*^26^. We also have shown *C. difficile* exacerbates distal colonic tumorigenesis in *Apc^Min/+^*mice^8^. Therefore, we interrogated the *C. difficile*-induced transcriptional network activation in vitro using purified recombinant TcdB and normal human colonic organoids (colonoids). We incubated colonoids with sub-lethal concentrations (< 1 pM) of TcdB for 5 weeks (Figure 7A-B). TcdB exposure led to multi-layered colonoids containing pleomorphic cells with variable nuclear-cytoplasmic ratios, similar to dysplastic crypts in tissue (Figure 7C). After a 2-week washout of toxin, we performed single-cell RNA-sequencing of the colonoids. Differential gene expression analysis of the TcdB-exposed colonoids revealed an increase in the WNT signaling pathway as evident by significant upregulation of *CTNNB1*, *CCND1*, *CCND2*, and *OLFM4* (Figure 7D). We repeated the TcdB exposure in colonoids from a different individual and detected increased cytoplasmic β-catenin protein compared to vehicle controls (Figure 7E-G). GSEA using the Wikipathways gene lists showed enrichment of YAP/TAZ signaling, proteoglycans in cancer, and microRNAs in cancer with TcdB-exposure (Figure 7H). Using the top 100 genes upregulated in TcdB-exposed colonoids, we performed transcriptional regulatory network analysis as we had done above for bacteria-gene expression correlations. Remarkably, the ELF3 transcription factor was predicted to be active in TcdB-exposed colonoids but was not significantly upregulated transcriptionally by TcdB (Figure 7I). ELF3 has previously been shown to increase β-catenin transcription with a subsequent increase in WNT pathway activation^52^. Finally, we examined ELF3 protein expression and demonstrated a significantly higher percentage of ELF3 nuclear localization in TcdB-exposed colonoids compared with vehicle control (Figure 7J-K). Based on the lack of *ELF3* transcriptional upregulation, we posit that TcdB is involved in the post-transcriptional regulation of ELF3 nuclear localization.

**Figure 7.**
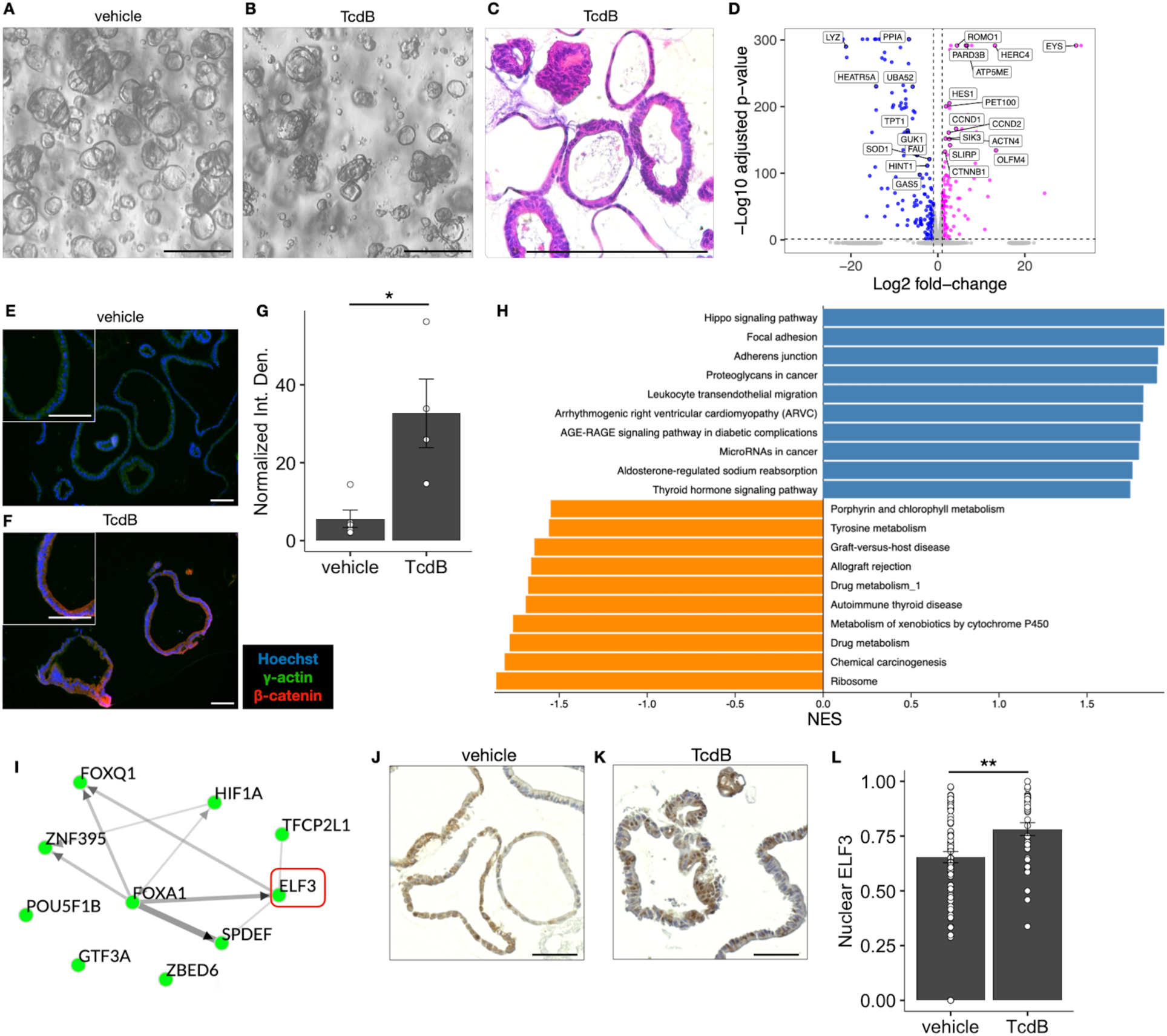
*C. difficile* toxin B (TcdB) induces activation of ELF3 and upregulation of β-catenin. (*A*) Phase contrast microscopy images of normal human colonic organoids (colonoids). (*B*) Colonoids incubated with sub-lethal concentrations (<1 pM) of TcdB for five weeks; scale bars = 200 μm. (*C*) Hematoxylin and eosin (H&E)-stained colonoids exposed to TcdB from (*B*). (*D*) Volcano plot showing differential gene expression analysis for TcdB-exposed colonoids (upregulated with TcdB in magenta on right side; downregulated with TcdB in blue on left side); *p*-values obtained with Wilcoxon rank-sum test. (*E-F*) Immunofluorescence of colonoids showing β-catenin upregulation at the protein level (red) in TcdB-exposed colonoids; scale bars = 100 μm; n = 4 technical replicates/group. (*G*) Normalized integrated density of β-catenin immunofluorescent staining from images in (*E-F*); *p*-value obtained with Wilcoxon rank-sum test. (*H*) Bar plot showing GSEA results using WebGestalt^63^. TcdB-exposed colonoids are enriched for the Wikipathways listed next to the blue bars and depleted for the pathways next to orange bars. NES = Normalized Enrichment Score; all scores had a False Discovery Rate < 0.05. (*I*) ChEA3-derived transcription factor regulatory network predicted by the upregulated genes from TcdB-exposed colonoids in (*E*). (*J-K*) Immunohistochemistry staining of ELF 3 in human colonoids exposed to vehicle or TcdB; scale bars = 100 μm. (*L*) Quantification of nuclear ELF3 staining from the images in (*J-K*); *p-*value obtained with Wilcoxon rank-sum test; \**p* < 0.05, \*\**p* < 0.01. n = 33-78 organoids/group.

Together, we have illustrated how the deep characterization of bacterial niches in CRC with spatial resolution can uncover functional relationships between low-abundance species and tumor cell transcriptional programming.

## Discussion

This CRC atlas uniquely characterizes the TME with an emphasis on defining intratumoral bacterial niches (Figure 5H). Although there are reports describing bacterial colocalization and CRC biological features^17,19,53^, the biogeographical association between microbes and the TME has been understudied. Most microbiome investigations related to CRC are limited to the quantification of microbial taxa from feces or bulk tumors without providing spatial resolution of intratumoral niches and the associated tumor transcriptomic profiles^11,54^. The low relative abundance and stability of bacterial mRNA compared with host transcripts, poor permeabilization of bacterial membranes, and lack of bacterial mRNA polyadenylation limit the ability to incorporate bacteria into TME studies. A benefit of using GeoMx was incorporating custom probes targeting bacteria previously associated with CRC as well as commensals (Figure 2E). Uniquely, we validated custom probes that queried Gram-positive species abundance in CRC.

Another unique aspect of our study was the selection of an optimal cohort. At the time of surgical resection, the clinical standard-of-care did not include preoperative oral antibiotics. Currently, CRC resections are performed after oral antibiotics and intravenous antibiotics, because these measures have been shown to reduce anastomotic leakage and postoperative infections^55,56^. Our cohort is limited by its small sample size, and there may be differences in CRC epidemiology between Malaysia and the United States. However, we previously demonstrated the microbiota and BFs show identical features between Malaysia and United States populations^21^. While an individual’s route to CRC may differ, the host-microbe associations may be more similar than different by the time of CRC diagnosis.

Our findings suggest ROIs with low bacterial burden had a vigorous immune response (Figure 2C). These results contrast published data associating stronger immune responses with regions of high bacterial burden, not low^20^. Both outcomes are feasible and may depend on the bacteria species present or type of ROI within the tissue. For example, we largely prioritized our ROI selection based on transitional features between normal and non-normal histopathology. Although our 16S data analysis suggested prominent Fusobacteria in multiple samples, *Fusobacterium* spp. were not as prevalent in the non-necrotic GeoMx areas. This results likely relates to *Fusobacterium* not being significantly elevated in Stage I CRC but developing blooms with later stages^24^. In contrast, Galeano Niño *et al*. selected ROIs based on bacterial burden rather than transition zones between histopathological subtypes, which may have led to preferential selection of necrotic regions with abundant Fusobacteria. We manually curated ROIs that preserved the near-necrotic host tumor cells (Supplementary Figure 6). The enrichment of epithelial-to-mesenchymal transition and coagulation genes in necrotic areas is consistent with Adrover et al. who mechanistically associated necrosis with tumor progression^37^. Additionally, we detected differences in interferon responses across different bacterial niches (Figure 5H), supporting the hypothesis that different microbial communities are associated with locally impactful tumor immune microenvironments that may foster immune stimulation versus suppression.

Our prior precancer atlas defined the differences between conventional adenoma-to-CRC progression and sessile serrated adenoma progression from metaplastic transition to MSI^high^ CRC^13^. Here, we focused on stage I and II CRC to capture intratumoral transition zones between histopathological features in a treatment-naïve cohort. BF status was only apparent from FISH and not by the composition or diversity measured by bulk 16S alone. Therefore, BF status largely represents a biogeographical shift in bacterial abundance due to mucus invasion of the associated luminal bacteria, as opposed to a compositional shift, consistent with our prior studies^21,22,27^.

To our knowledge, prior CRC studies have not included targeted analysis of Gram-positive species^5,19,20^. The Gram-positive *E. faecalis*, *P. micra*, *P. stomatis*, and *R. gnavus* were 4/5 of the most abundant species in the bacterial niches (Figure 3A). This representation was also observed in the unbiased 16S data (Figure 2C). The high abundance of Gram-positive species in our cohort might be a consequence of the lack of preoperative oral antibiotics. The relative abundance of *E. faecalis* in Niche 6 was unique. Also, *Streptococcus gallolyticus* was found in both 16S and custom probe detection in GeoMx ROIs deep within the tumors (Figure 3E). *S. gallolyticus* (formerly *S. bovis*) bacteremia and endocarditis have long been associated with CRC^57,58^.

Perhaps the most unexpected discovery from this atlas was the disproportionate impact of the low-abundance organisms, like *C. difficile* and *E. aldenensis*, on host gene expression (Figure 5F, Figure 6A). We validated the role of *C. difficile* TcdB in promoting the nuclear translocation of ELF3, cytoplasmic accumulation of β-catenin, and upregulation of WNT target genes (Figure 7). Others have shown low-abundance species contribute to critical molecular functions for the gut microbiome-host relationship, such as bacterial pilus assembly and methane production^59^. Metagenomics data have suggested about 45% of microbial genes with potentially critical functions in the gastrointestinal ecosystem are present in less than 10% of bacterial genomes^60^. Commensal species, like *M. morganii*, were recently discovered to produce genotoxic metabolites that exacerbate CRC in mouse models^61^. Our own work has demonstrated how human biofilm-associated *C. difficile* accelerates distal colonic tumor formation in mouse models and potentially human CRC^25,26^, despite making up approximately 1% or less of the relative abundance of the colonic microbiome^8^. Even during symptomatic *C. difficile* infection, its average abundance is less than 2%^62^. Our findings support the growing appreciation for low-abundance organisms with important functions in gut homeostasis and disease.

In summary, we created a multi-omic atlas of CRC with bacterial niches of both naturally occurring, biogeogrpaphic origins as well as niches identified by computational clustering. We deeply characterized these niches at the level of each individual with CRC, tissue, cell type, and transcriptomic program. The intriguing relationships we uncovered provide a foundation for future mechanistic studies. We hypothesize that a comprehensive understanding of bacteria-tumor interactions will come from studies focused on low-abundance organisms, like *C. difficile*, that drive functional changes in cancer biology.

## Abbreviations

AC: adenocarcinoma
BMI: body mass index
CODEX: co-detection by indexing
CMS: consensus molecular subtyping
CRC: colorectal cancer
BF: biofilm
DSP: digital spatial profiler
FFPE: formalin-fixed paraffin-embedded
FISH: fluorescence *in situ* hybridization
GSEA: gene set enrichment analysis
H&E: hematoxylin and eosin
HGD: high-grade dysplasia
MSI: microsatellite-instable
MSS: microsatellite-stable
OTU: operational taxonomic unit
PanCK: pan-cytokeratin
ROI: region of interest
tSNE: t-distributed stochastic neighbor embedding
WTA: whole transcriptome atlas

## Author Contributions

NOM, JLD, DBL, MJS, KSL, CLS, and RJC designed and funded the research. NOM, JLD, EHG, WEB, HML, AJS, HK, JQ, ALG, XL, ZM, FR, and JC performed research. MP, BRH, ES, MR-S, AW, MKW, CC, SM, LD, QL, KSL, RJC, and CLS analyzed data. JWW, TI, ACR, and JV assisted in the setup, recruitment, or handling of clinical specimens. NOM, JLD, and RJC wrote the paper with input from all authors.

## Data Transparency Statement

All raw and processed data are openly available within the HTAN data repository (https://data.humantumoratlas.org/data-access). All code for analysis and visualizations are available from the authors upon request.

## Acknowledgements

We thank the participants, staff, and scientists who contributed to this study, especially the staff at HTAN member centers. We thank Dr. Emma Allen-Vercoe (University of Guelph) for generously providing 3 bacterial strains for our validation array: *Fusobacterium periodonticum* (EAV CRC 1/1/44), *Gemella haemolysans* (EAV CRC CC7/3 CNA4), *Peptostreptococcus stomatis* (EAV CRC CC3/4 C14). We thank James R. White for aiding with 16S rRNA amplicon sequencing visualizations. We thank Sarah E. Glass for manuscript editing. The following core facilities at Vanderbilt University Medical Center performed experiments or data analysis: Tissue Pathology Shared Resource, VANTAGE center for advanced genomics, and Digital Histology Shared Resource.

---

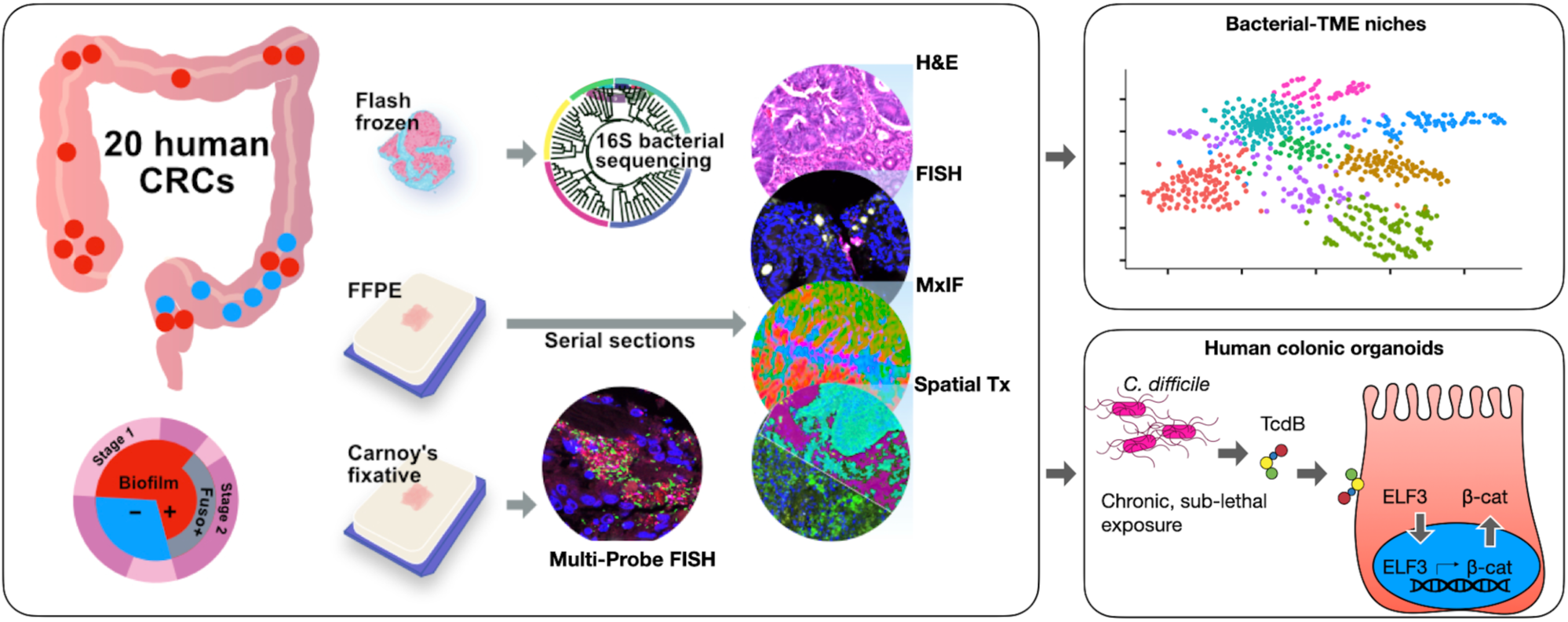

## Supplementary Materials

**Supplementary Table 1**: Malaysia nested cohort patient metadata. HTAN = Human Tumor Atlas Network. BMI = Body-Mass Index.

**Supplementary Table 2**: Bacterial-Tissue Microarray (Bac-TMA) details including the sequences of custom bacterial probes and how they were pooled, the summaries of the probe validation, and a list of the 30 isolates from the 3728T consortium^8^.

**Supplementary Table 3**: Bacterial isolates and culture conditions. ANA = anaerobic. AER = aerobic. BHI = brain-heart infusion. BRU = Brucella blood agar.

**Supplementary Table 4**: CODEX Samples, antibodies, and cycle information

**Supplementary Table 5**: Niche Summary. ns = not significant

**Supplementary Table 6**: Bacterial abundance correlations with tumor gene expression. FDR = False Discovery Rate.

**Supplementary Table 7**: Fluorescence in situ hybridization (FISH) Probes

**Supplementary Table 8**: qPCR primers and probes

**Supplementary Figure 1:**
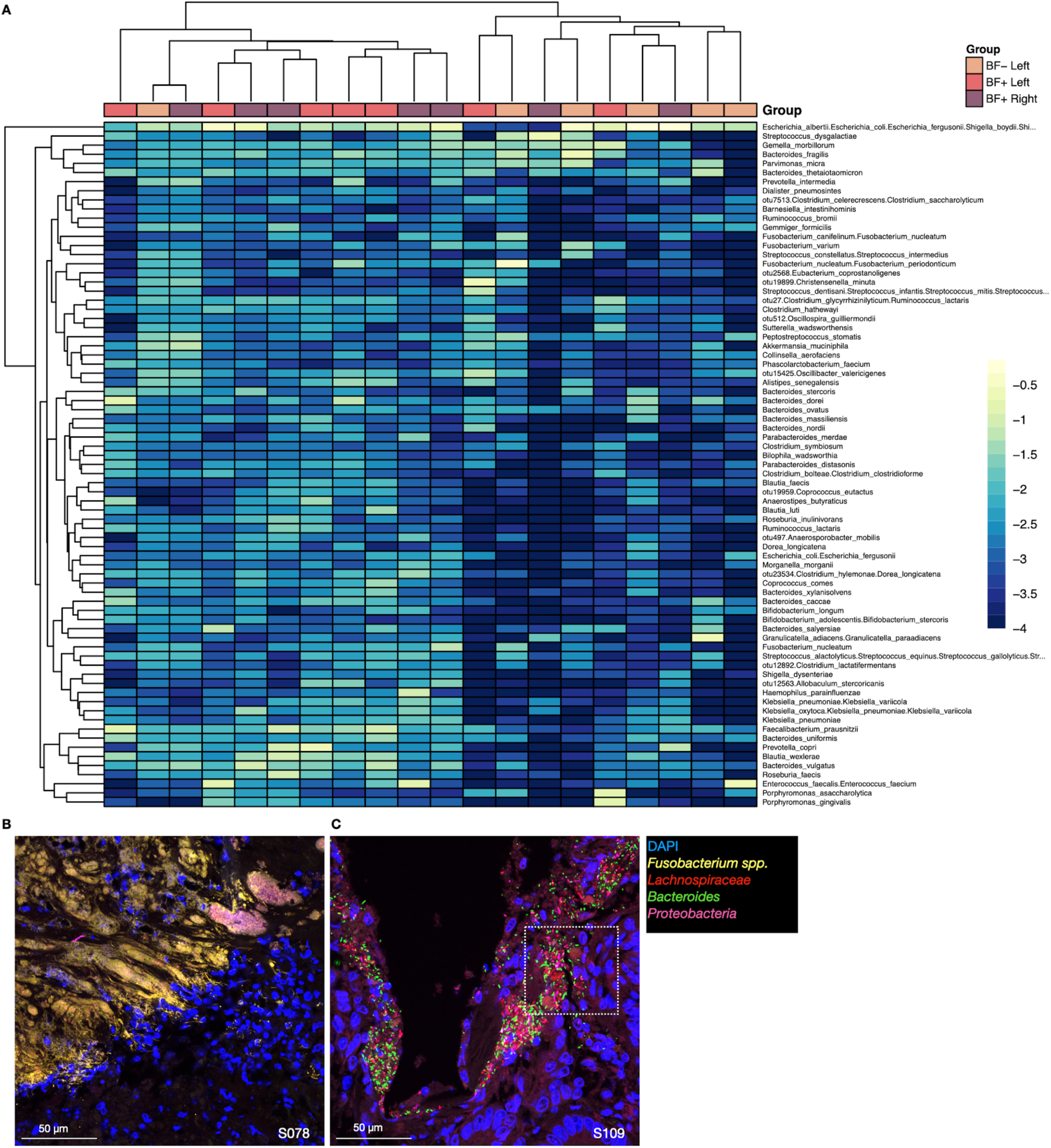
(*A*) Bulk 16S data displayed as a heatmap with colors representing normalized, log-transformed counts and grouping by BF status with colon sidedness (Left versus Right). (*B-C*) Multi-probe FISH microscopy demonstrating a *Fusobacterium* spp. bloom (*B*) and a polymicrobial BF (*C*). The white dotted box in image (*C*) corresponds to the magnified image in Figure 1F.

**Supplementary Figure 2:**
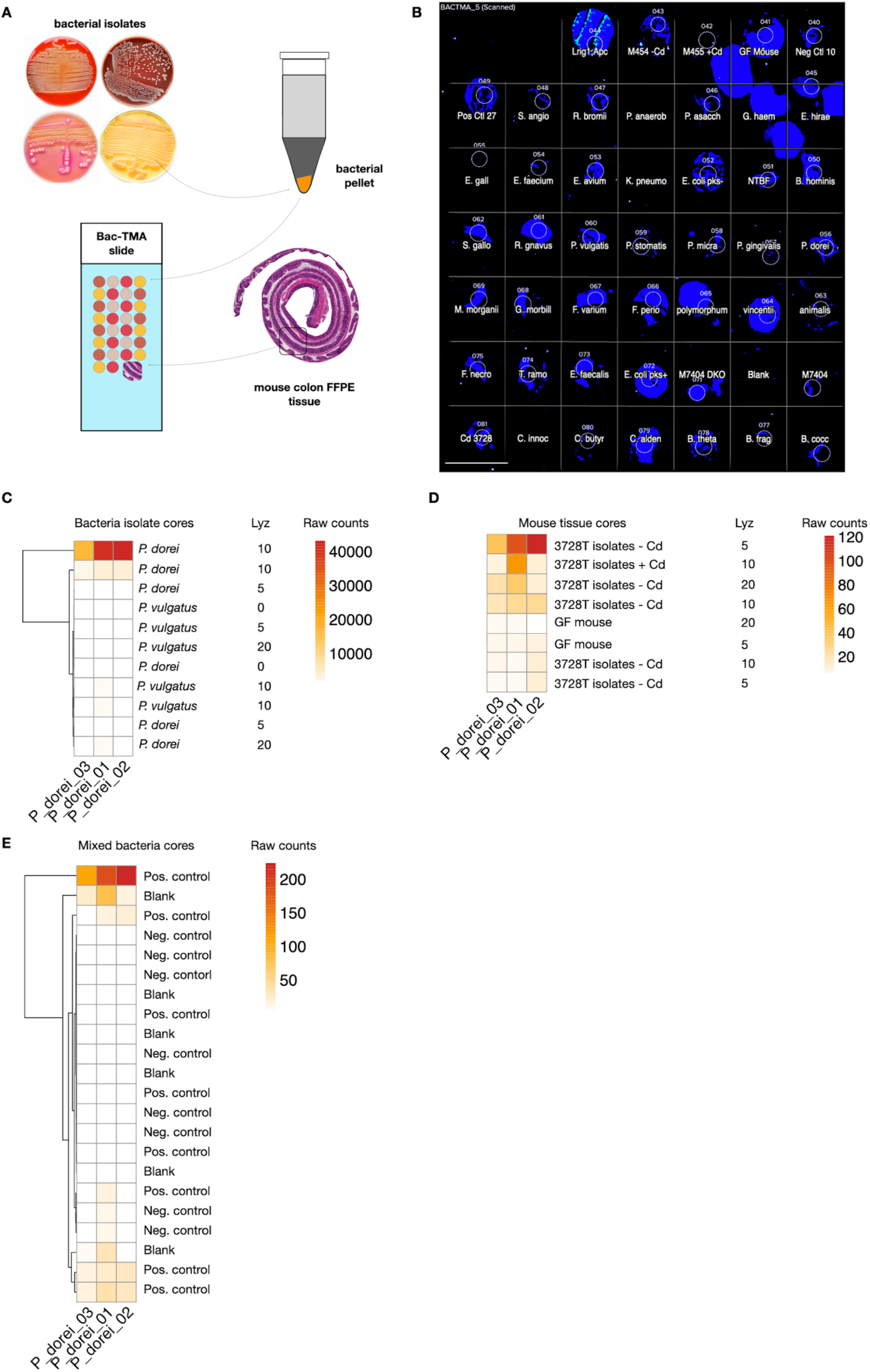
Custom Bacteria-Tissue Microarray (Bac-TMA) for experimental validation of species-specific GeoMx probes. (*A*) Schematic of the Bac-TMA creation. (*B*) Example of a fluorescence image from the GeoMx Digital Spatial Profiler of the Bac-TMA with nuclei in blue and ROIs represented by white circles; scale bar = 1 mm. (*C*) Heatmap of bacterial abundance raw counts for three different probes against *P. dorei*. The closely related *P. vulgatus* species was used as a negative control. Different lysozyme (Lyz) concentrations (mg/mL) were tested to define the optimal conditions for probe hybridization to both Gram-positive and Gram-negative bacteria. (*D*) The three *P. dorei* probes were tested against cores of mouse colonic tissue from animals containing the 3728T isolates^8^ plus or minus *C. difficile* (Cd) and negative control germ-free (GF) mouse colonic tissue. (*E*) Heatmap of *P. dorei* probes tested against cores of mixed bacterial pellets versus blank paraffin or negative control pellets that did not contain *P. dorei*.

**Supplementary Figure 3:**
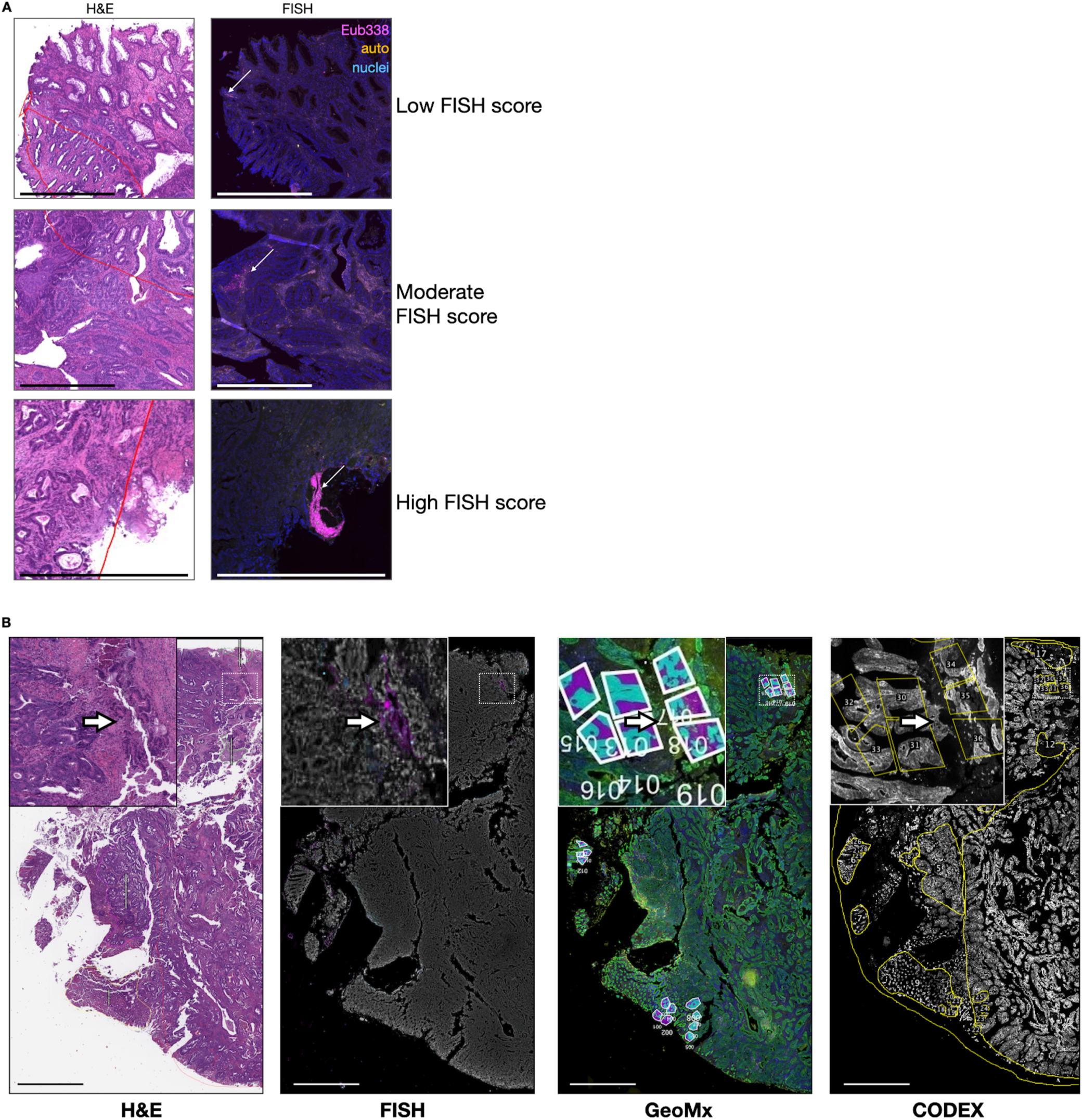
Image registration for identifying areas of high bacterial load and spatial transcriptomics ROIs. (*A*) Paired H&E and FISH microscopy images show examples of low, moderate, and high FISH scores that were used to annotate the GeoMx ROIs; scale bars = 1 mm. Red lines on the H&E images are from the pathologists’ annotations of the histopathological features. (*B*) Registered images from different modalities show an example of how ROIs were chosen on the GeoMx Digital Spatial Profiler (including segmented ROIs) and how annotations were made on the CODEX images (yellow lines); scale bars = 1 mm.

**Supplementary Figure 4:**
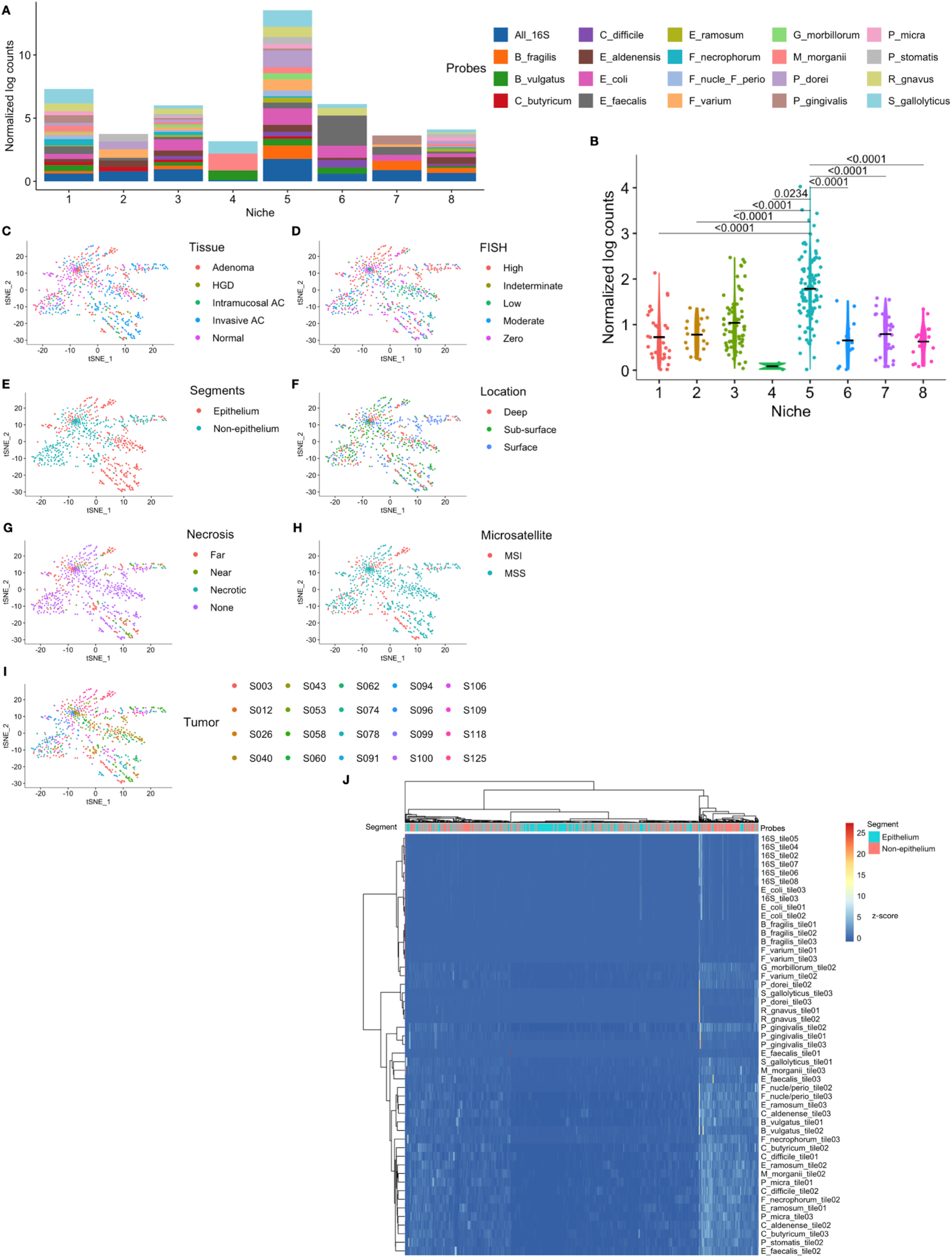
(*A*) Stacked bar plot showing the abundance and diversity of bacteria species within each niche. The same data are plotted as a heatmap in Figure 2F. (*B*) Violin plot illustrating the total bacterial abundance (All_16S) in each niche; p-values were obtained from Dunn’s test. (*C-I*) tSNE plots of the GeoMx ROIs using the same coordinates as in Figure 2D but colored based on different tumor features. (*J*) Heatmap of all custom bacterial probes (rows) and all GeoMx ROIs (columns) colored with a normalized z-score representing abundance. Hierarchical clustering shows modest clustering based on epithelium versus non-epithelium.

**Supplementary Figure 5:**
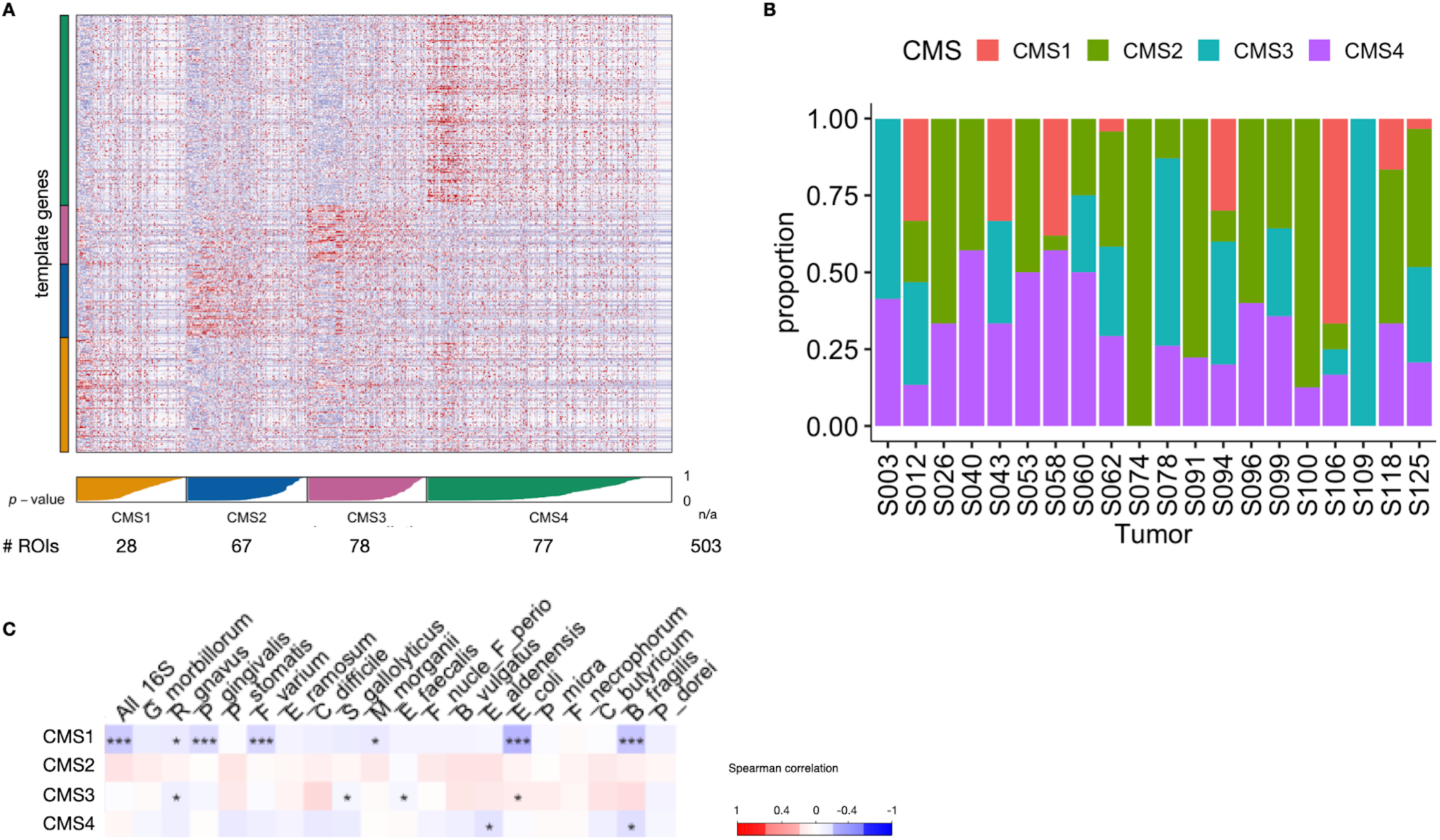
CMS predictions and correlations to bacterial detection. (*A*) Heatmap with CMS classification predictions for each GeoMx ROI. Notably, 503 ROIs did not have a statistically significant prediction. (*B*) Of the ROIs that had a calculable CMS classification, their proportions are shown as a stacked bar plot for each tumor, reflecting the overall heterogeneity of the samples in this cohort. (*C*) Correlation plot showing the relationship between specific bacterial abundance and the CMS classifications; \**p* < 0.05 and *** *p* <0.001.

**Supplementary Figure 6:**
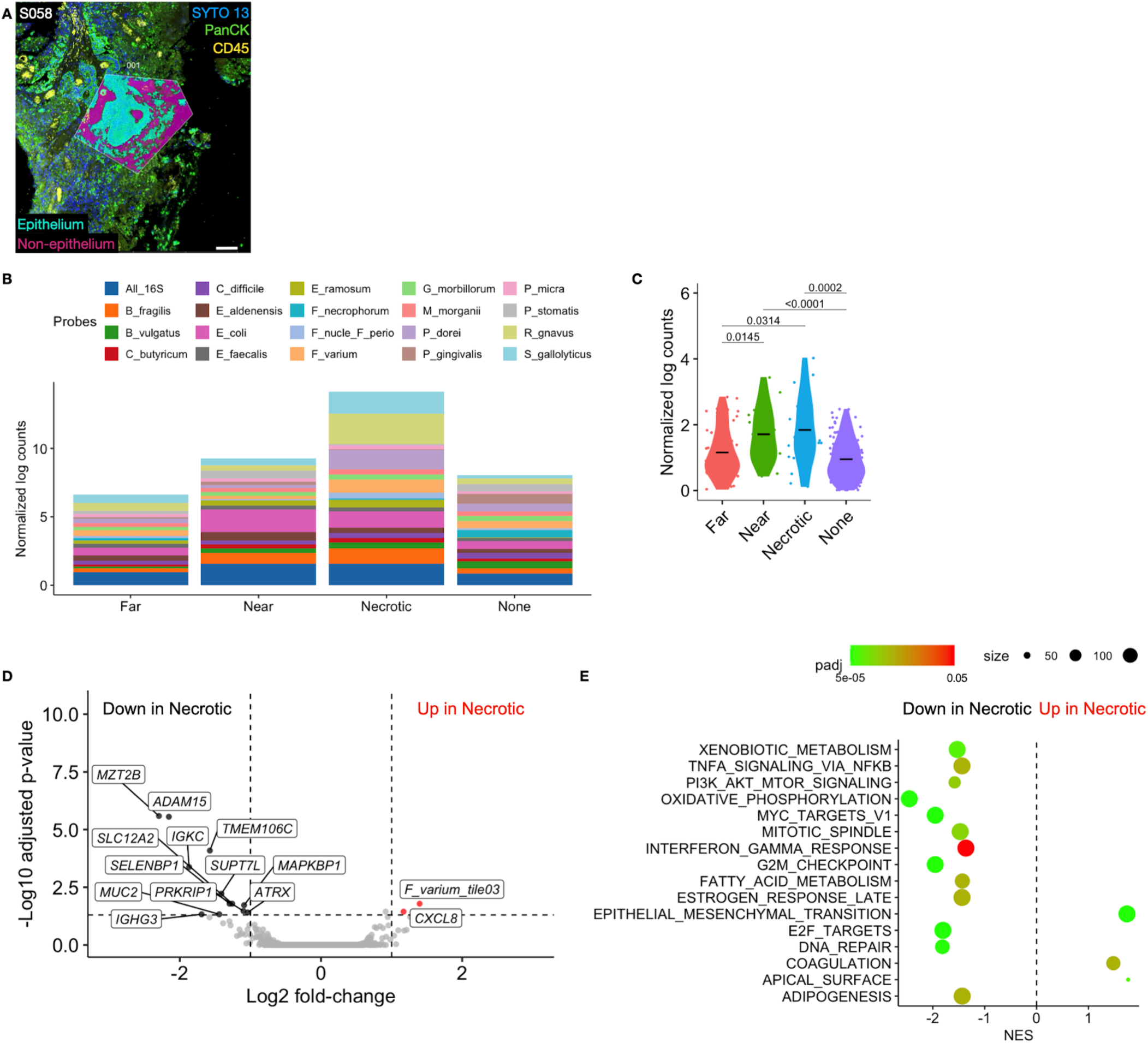
Total bacterial abundance was high in necrotic tumor regions. (*A*) Immunofluorescence microscopy example of a necrotic region within tumor S058. Epithelium (cyan) and non-epithelium (magenta) masks are shown in (*A*); scale bar = 100 μm. (*B*) Bacterial load was highest in GeoMx areas around necrosis. “Necrotic” regions were areas bordering necrosis (including luminal necrosis and ulcerated tumor surfaces), “Near” includes areas within 1 mm of necrosis, “Far” includes areas within a necrosis-bearing tumor but not within 1 mm, and “None” includes areas in tumors without any necrosis. (*C*) Total bacterial counts as detected by All_16S probes were significantly higher in areas bordering necrosis compared with areas far from or without any necrosis; *p*-values obtained from Dunn’s test. (*D*) Volcano plot showing differential gene expression analysis comparing necrotic areas to all other segments. (*E*) GSEA comparing necrotic to non-necrotic areas using the MsigDB Hallmarks database; Normalized Enrichment Score (NES).

**Supplementary Figure 7:**
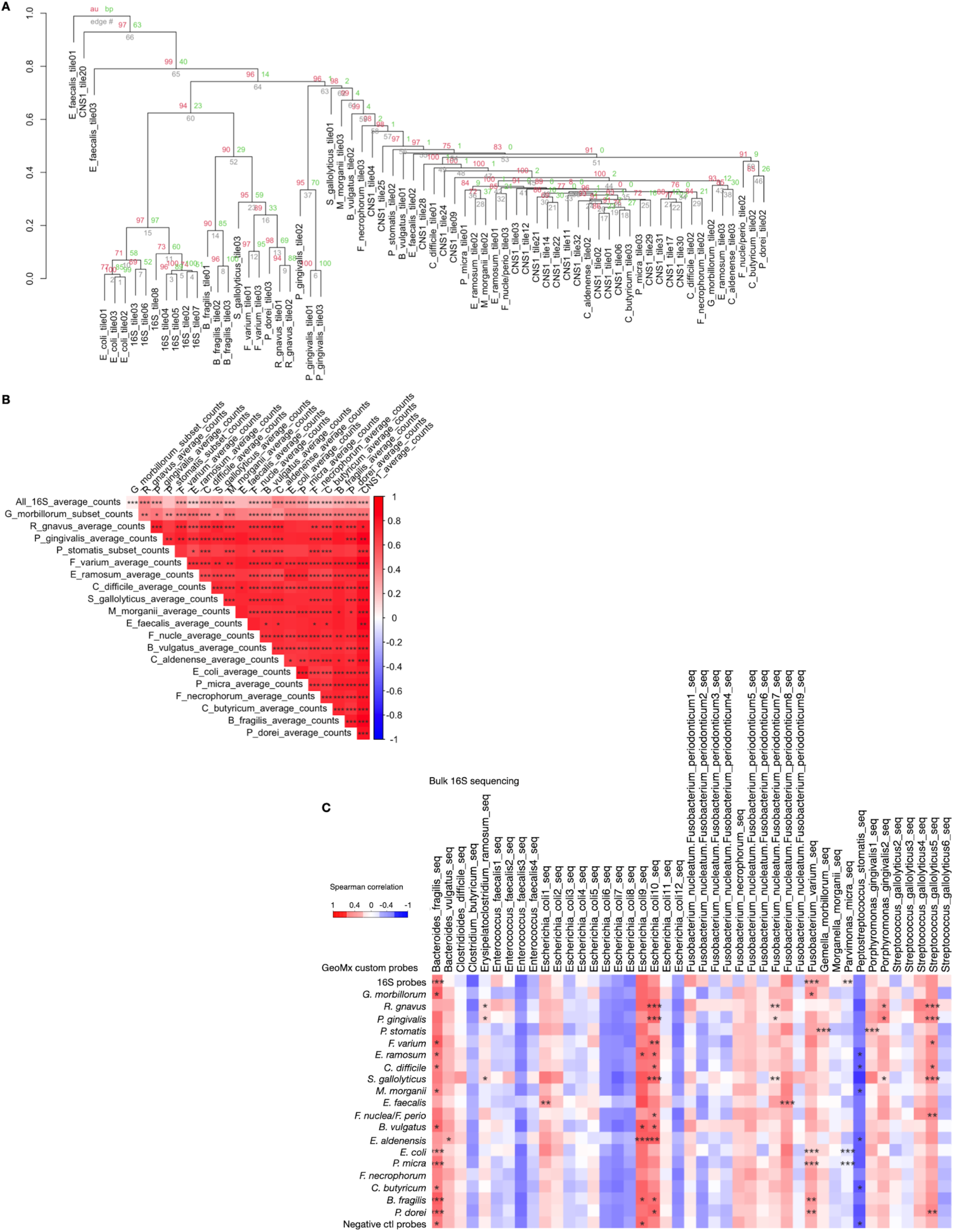
(*A*) Hierarchical clustering of the individual custom bacterial probes demonstrating similarity between probes designed against the same species, justifying our use of geometric means for each species. (*B*) Correlation plot for mean abundance of custom probes from the GeoMx ROIs. (*C*) Correlation plot showing the integration of bulk 16S data (20 most abundant OTUs) with custom bacterial species probe detection from GeoMx. \**p* < 0.05, \*\**p* < 0.01, and *** *p* <0.001.

**Supplementary Figure 8:**
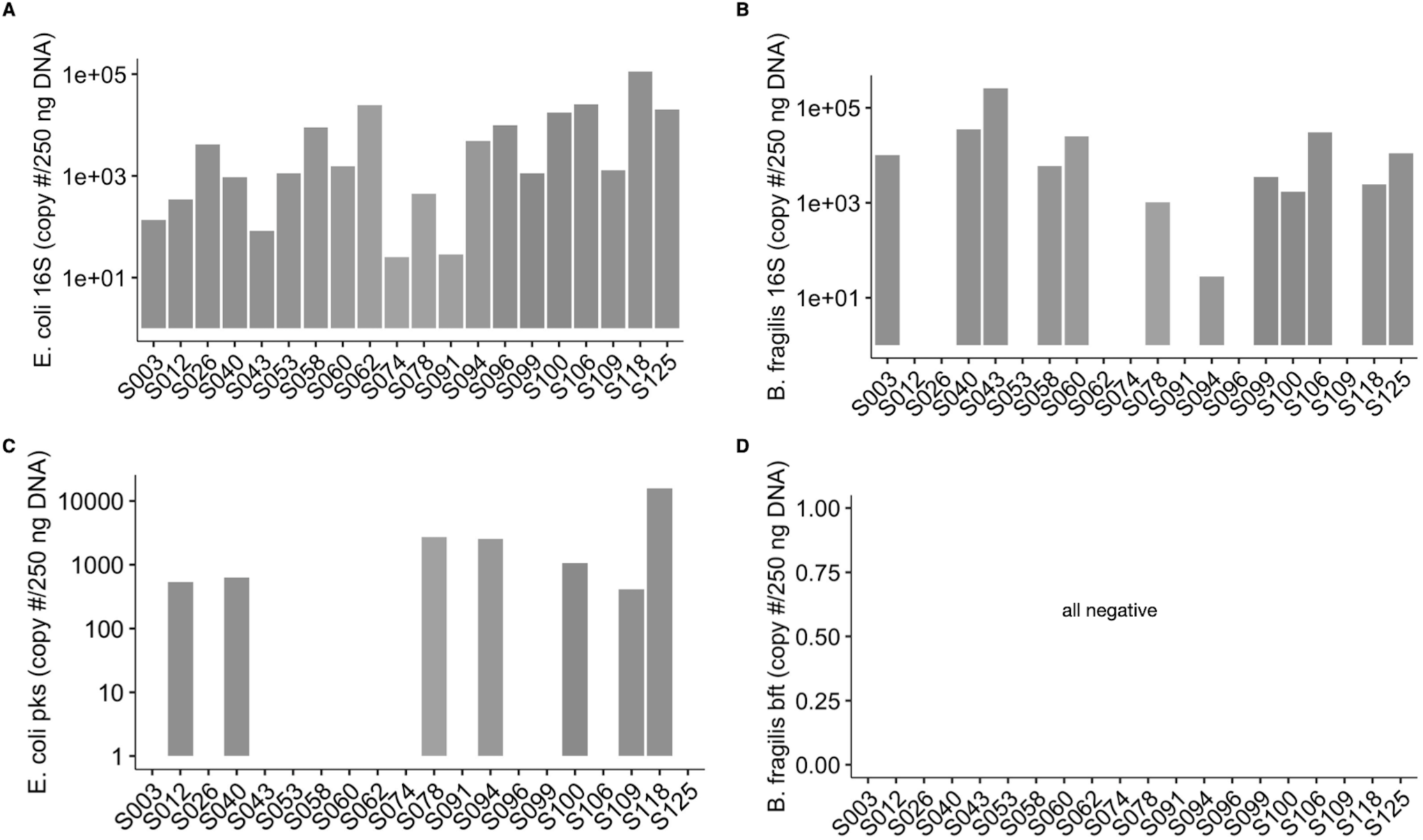
*E. coli* and *B. fragilis* abundance and toxin expression. (*A*) Total *E. coli* and (*B*) total *B. fragilis* 16S gene abundance by qRT-PCR from flash-frozen tumor specimens. (*C*) Abundance of polyketide synthase (pks)^+^ *E. coli* in each tumor as determined by qRT-PCR from flash-frozen tumor specimens. (*D*) The *Bacillus fragilis* toxin (*bft*) gene was not found in any of the tumors by qRT-PCR.

**Supplementary Figure 9:**
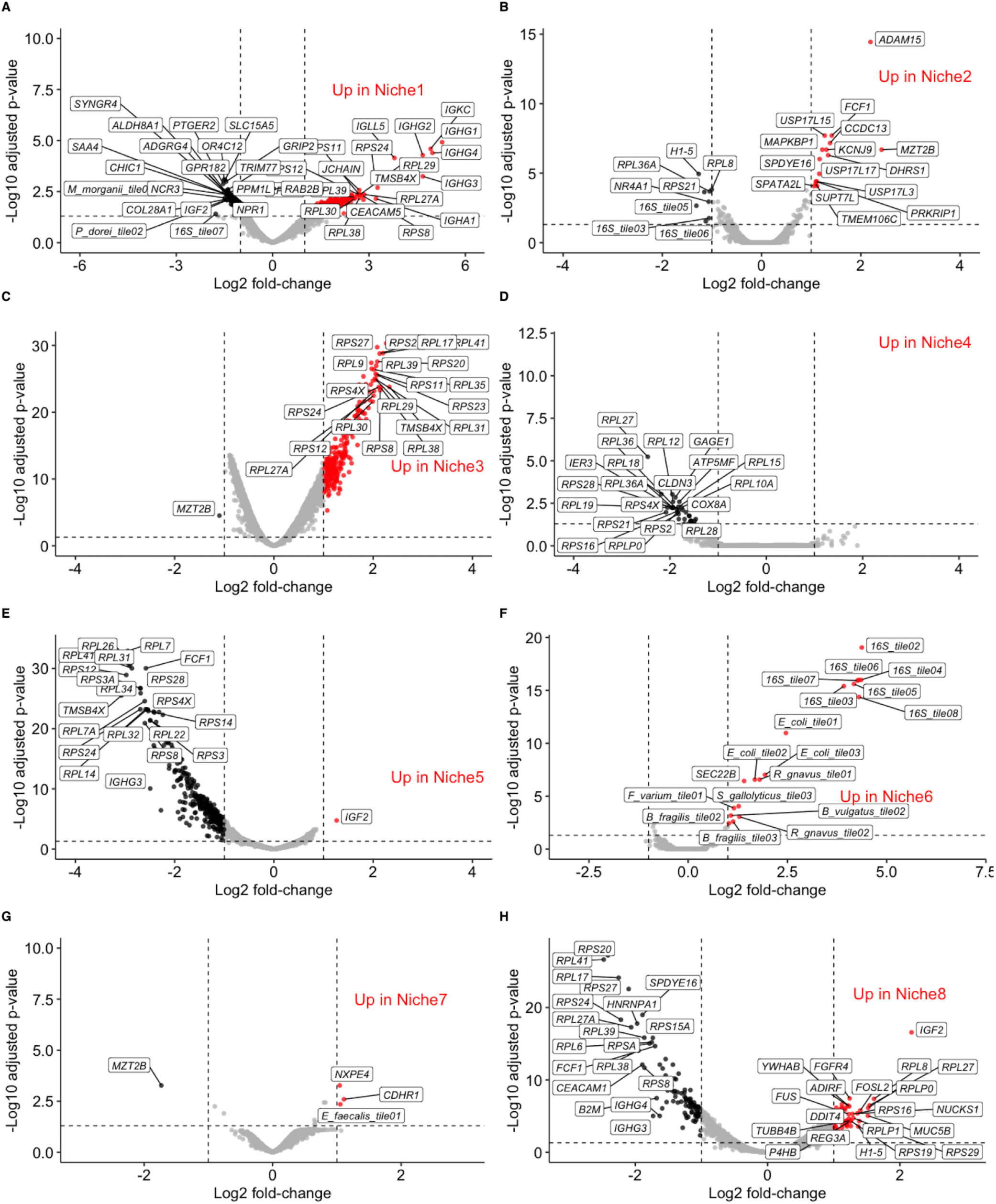
Differential gene expression in the niches. (*A-H*) Volcano plots showing the uniquely upregulated and downregulated genes in each niche compared to all the others using linear mixed modeling. Adjusted *p*-values were obtained using the Benjamini-Hochberg procedure.

**Supplementary Figure 10:**
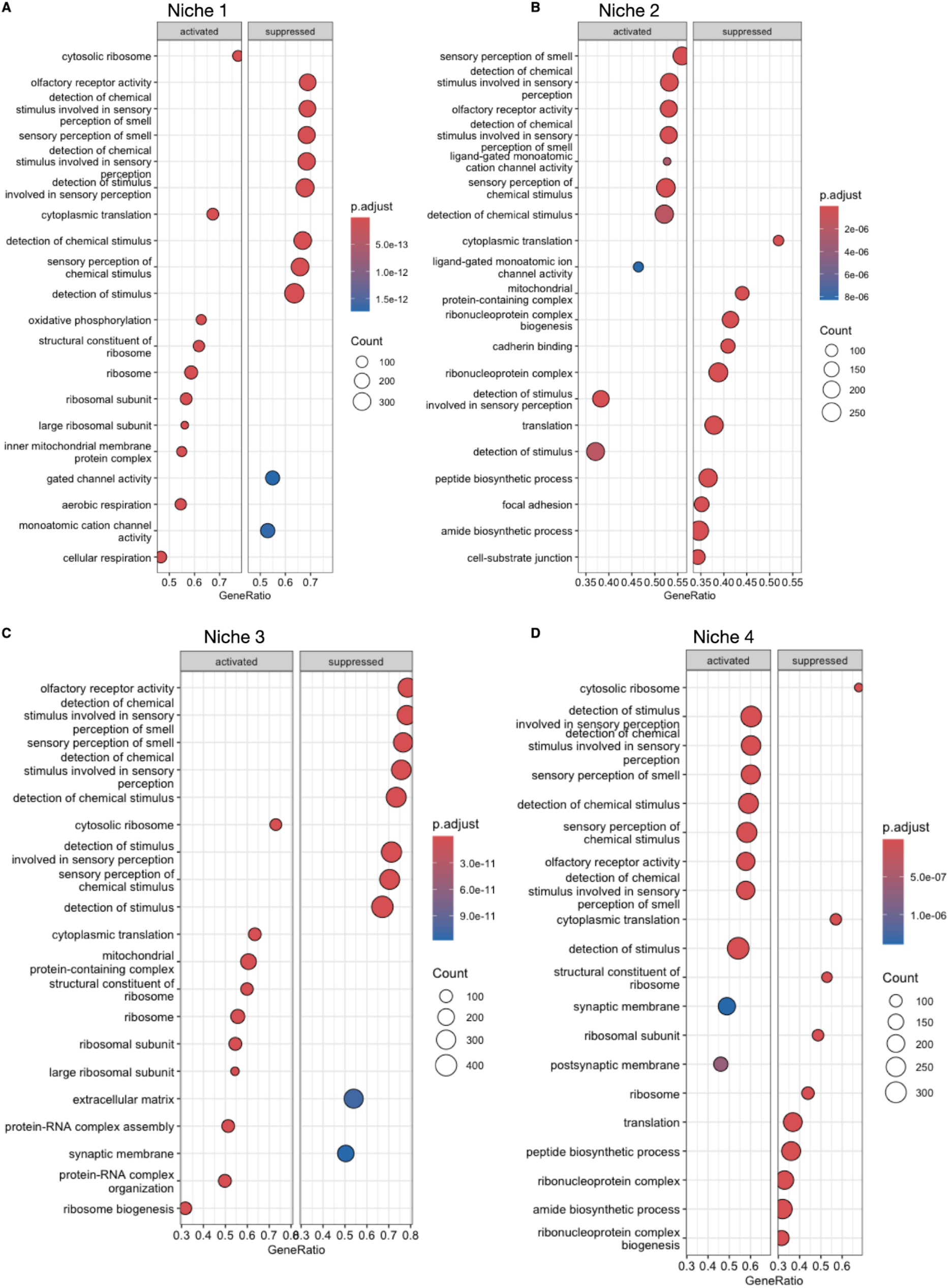
GSEA using gene ontology for niches 1-4. (*A*-*D*) GSEA plots showing the activated (enriched) and suppressed (depleted) pathways in each niche based on the differentially expressed genes in Supplementary Figure 9.

**Supplementary Figure 11:**
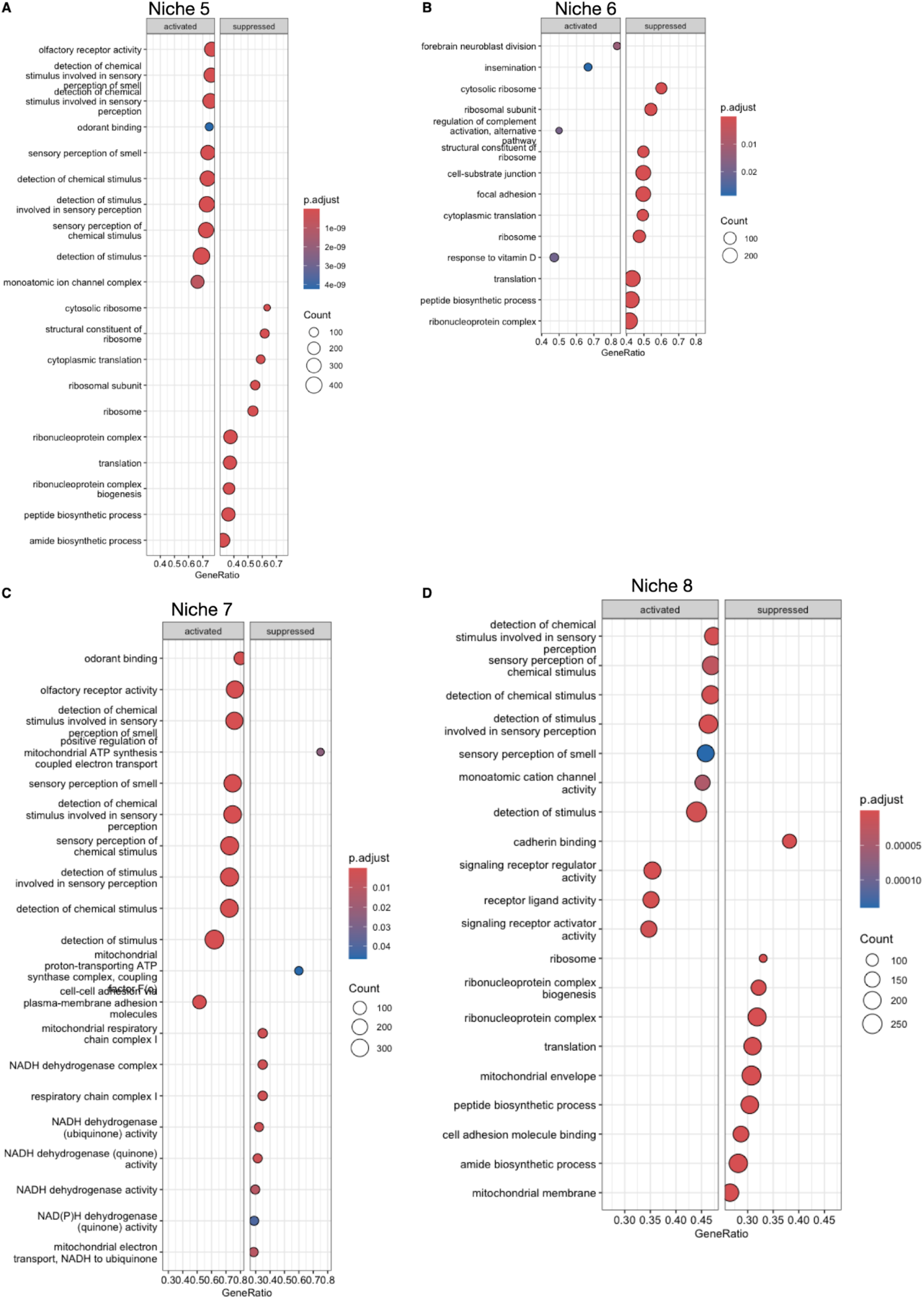
GSEA using gene ontology for niches 5-8. (*A*-*D*) GSEA plots showing the activated (enriched) and suppressed (depleted) pathways in each niche based on the differentially expressed genes in Supplementary Figure 9.

**Supplementary Figure 12:**
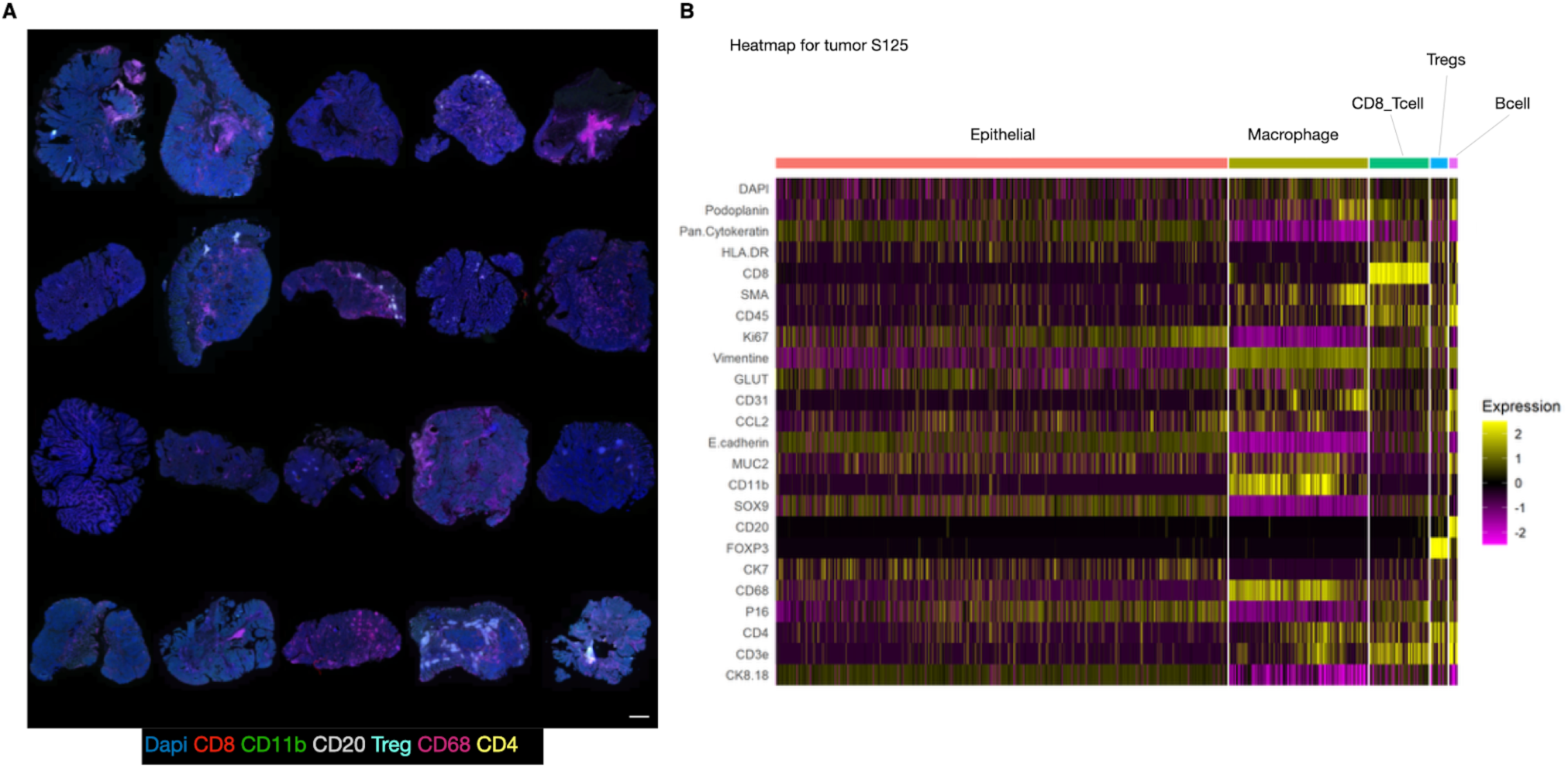
CODEX multiplex immunofluorescence. (*A*) Montage of all 20 tumor sections with pseudo-colored fluorescence channels chosen to reflect the immune infiltration in each specimen; scale bar = 1 mm. (*B*) Heatmap of normalized expression for the different CODEX markers (rows) in each of the cells (columns) segmented from immunofluorescence images. This example from tumor S125 shows how cell type assignments were given to specific clusters of cells.

**Supplementary Figure 13:**
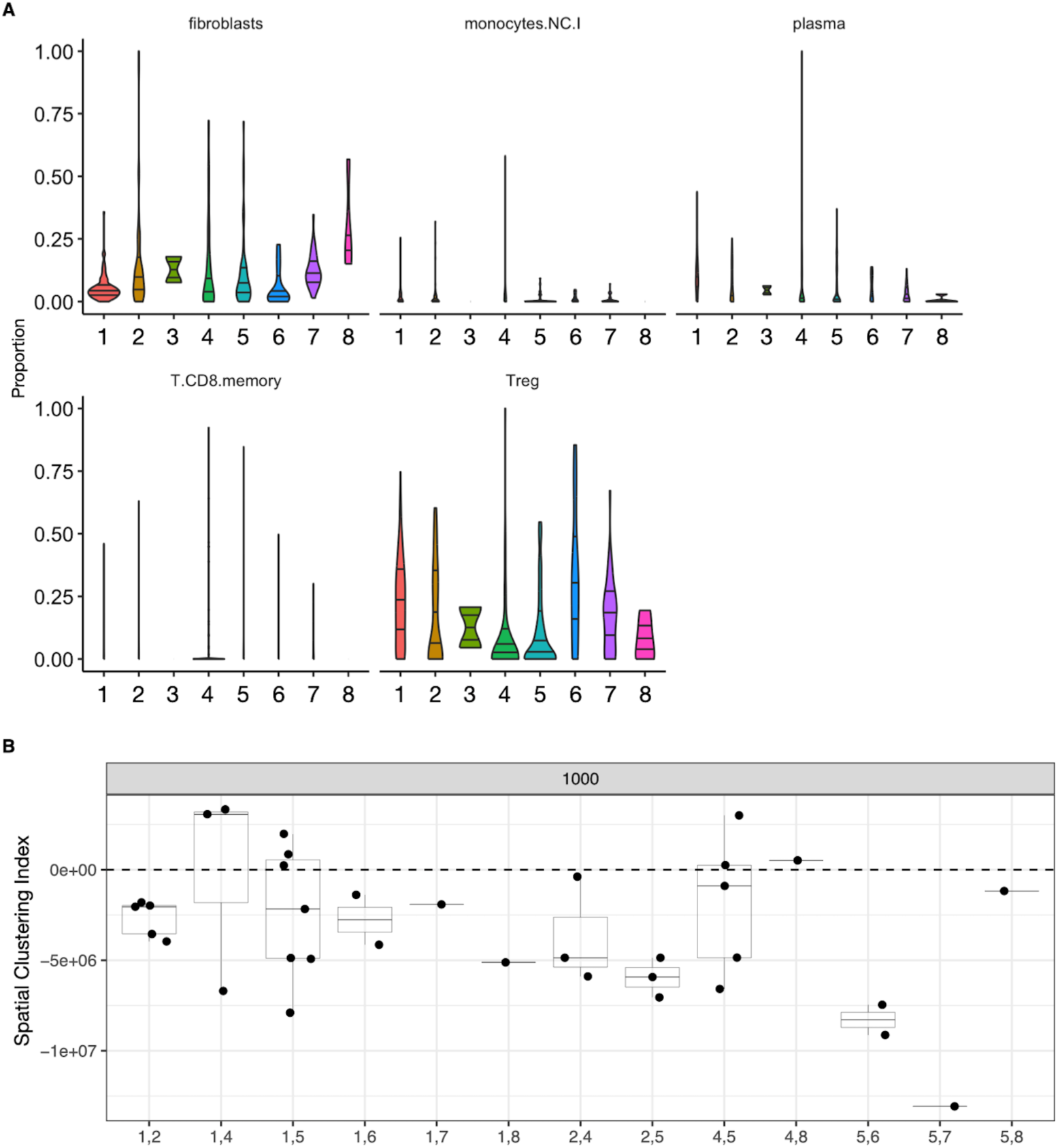
(*A*) Spatial deconvolution of GeoMx transcriptomic data for the non-epithelial segments. The *safeTME* dataset was used as a benchmark^65^. Only cell types with significant differences (FDR < 0.05) between niches are shown. (*B*) Spatial clustering index for pairs of niches that appear in the same tumors. The medians of these pairs generally fall below zero, which indicates that the listed pairs of niches do not co-localize with each other.

**Supplementary Figure 14:**
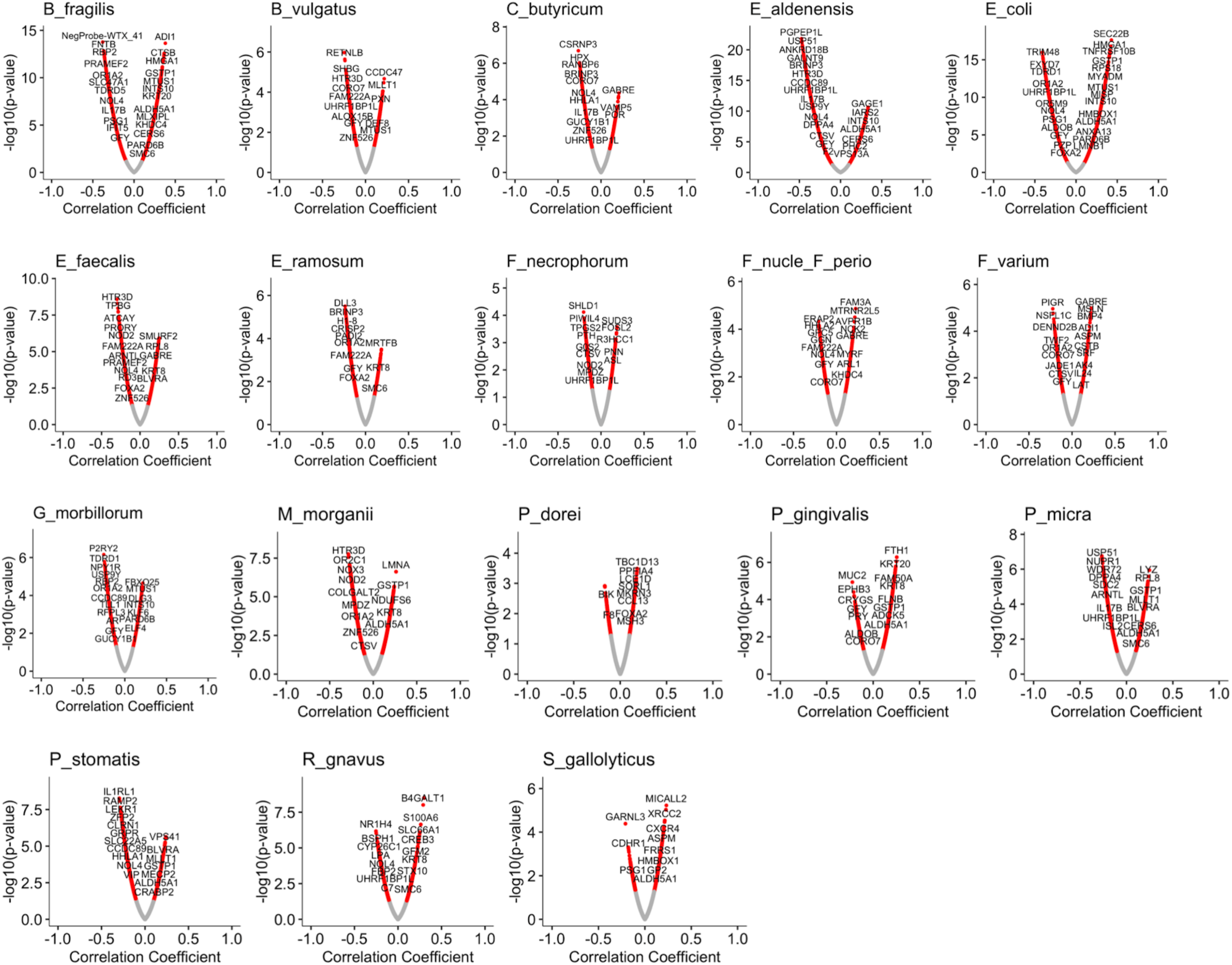
Correlations between bacterial species abundance and tumor gene expression. Each panel shows a different volcano plot of the specific genes with statistically significant correlations between their expression and the abundance of the listed bacterial species. Gray is non-significant; red indicates significant genes.

**Supplementary Figure 15:**
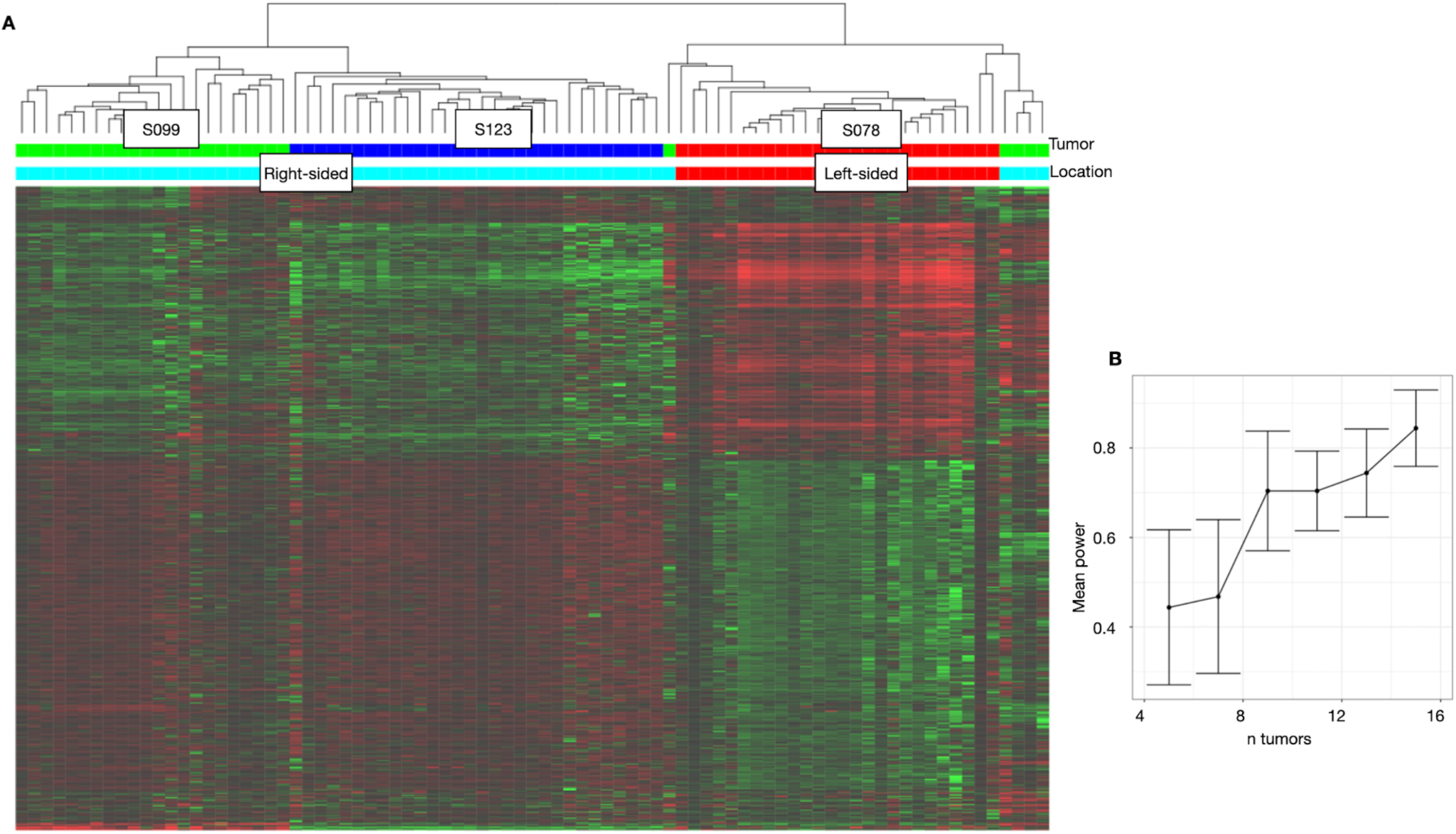
(*A*) Heatmap for pilot testing of GeoMx with three tumors. Genes are rows, and ROIs are columns. (*B*) Power analysis using preliminary data from (*A*) shows that we would have 80% power to achieve statistically significant differences in spatial transcriptomics from 15-16 tumors.

